# Paradoxical Aortic Stiffening and Subsequent Cardiac Dysfunction in Hutchinson-Gilford Progeria Syndrome

**DOI:** 10.1101/790477

**Authors:** S-I. Murtada, Y. Kawamura, A.W. Caulk, H. Amadzadeh, N. Mikush, K. Zimmerman, D. Kavanagh, D. Weiss, M. Latorre, Z.W. Zhang, G.S. Shadel, D.T. Braddock, J.D. Humphrey

## Abstract

Hutchinson-Gilford Progeria Syndrome (HGPS) is an ultra-rare disorder with devastating sequelae resulting in early death, presently believed to stem primarily from heart failure secondary to central arterial stiffening. We analyze novel longitudinal cardiovascular data from a mouse model of HGPS (*Lmna*^*G609G/G609G*^) using allometric scaling and advanced computational modelling and show that a late-stage increase in pulse wave velocity, with associated diastolic dysfunction but preserved systolic function, emerges with a loss of aortic function, independent of sex. Specifically, there is a dramatic late-stage loss of smooth muscle function and cells and an excessive accumulation of proteoglycans along the entire aorta, which result in a loss of biomechanical function (contractility and elastic energy storage) and marked structural stiffening despite a distinctly low intrinsic material stiffness that is consistent with the lack of functional lamin A. Importantly, vascular function appears to be normal within the low stress environment of development, only to succumb progressively to pressure-related effects of the lamin A mutation and become extreme in the peri-morbid period. Because the dramatic life-threatening aortic phenotype manifests during the last quarter of life there may be a therapeutic window in maturity that could alleviate concerns with therapies administered during early periods of arterial development.

**Disclosures:** D.T.B is an equity holder in, and receives research and consulting support from, Inozyme Pharma, Inc. for therapeutics for ENPP1 deficiency. None of the other authors declare any conflict, financial or otherwise.

## INTRODUCTION

Hutchinson-Gilford Progeria Syndrome (HGPS) is a genetic disorder characterized by premature ageing with devastating consequences to cardiovascular and musculoskeletal tissues. Accelerated atherosclerosis in muscular arteries (Olive et al., 2010), which can lead to myocardial infarction or stroke, was thought to cause death in the early teens, but statins and lipid-lowering agents did not improve lifespan. HGPS also accelerates stiffening of central arteries, as inferred from pulse wave velocity (PWV), and thereby contributes to elevated central blood pressures (Gerhard-Herman et al., 2012). Increased central artery stiffness is an initiator and indicator of diverse cardiovascular diseases in the general population and thus a predictor of all-cause mortality (Vlachopoulos et al., 2010), particularly due to myocardial infarction, stroke, and heart failure (Mitchell et al., 2010). Importantly, a recent study revealed that left ventricular diastolic dysfunction was the most prevalent abnormality in HGPS (Prakash et al., 2018), and a clinical trial identified heart failure as the primary cause of death (Gordon et al., 2018). There is, therefore, a pressing need to investigate together the changes in central vessels and cardiac function. Given the scarcity of human data, mouse models enable more detailed study of both the underlying mechanisms and resulting clinical phenotypes.

HGPS arises from point mutations in the gene (*LMNA*) that encodes the cell nuclear envelope protein lamin A (Eriksson et al., 2003), which normally contributes to nuclear stiffness and transcriptional regulation (Verstraeten et al., 2008) and is highly mechanosensitive (Swift et al., 2013). Mutations can lead to an altered lamin A precursor, resulting in a truncated form of lamin A called progerin (from the Greek, *pro* [before] *geras* [old age]). Using a bacterial artificial chromosome approach, a transgenic mouse model was generated with the human mutation (c.1824C>T;pG608G). Central arteries from these mice exhibit fewer smooth muscle cells (SMCs), medial calcification, accumulated proteoglycans, disorganized collagen, and some fragmented elastic fibres (Varga et al., 2006). Such changes in composition and microstructure would be expected to compromise biomechanical functionality of the arterial wall. Indeed, it was suggested that arterial SMCs in this mouse “are especially vulnerable to the mechanical stress imposed on them” despite blood pressure remaining nearly normal (Capell et al., 2008). Surprisingly, detailed quantification of arterial function and properties remains wanting, though a recent study reported stiffening of the thoracic aorta and mesenteric artery in another mouse model of HGPS (del Campo et al., 2019), generated using a knock-in mutant allele that carries a c.1827C>T;p.G609G mutation (Osorio et al., 2011), denoted *Lmna*^*G609G/G609G*^. The associated cardiovascular phenotype has yet to be examined in detail, but there is loss of SMCs and calcification of the aortic media in *Lmna*^*G609G/+*^ mice (Villa-Bellosta et al., 2013) and decreased SMCs, decreased elastic fibre waviness, and increased collagen in *Lmna*^*G609G/G609G*^ mice (del Campo et al., 2019).

Normal ageing progressively and differentially affects the aorta along its length (Rogers et al., 2001; Ferruzzi et al., 2018), thus effects of HGPS should be quantified as a function of age and aortic location. Our aim was to biomechanically phenotype the entire aorta in adult male and female *Lmna*^*G609G/G609G*^ mice and to evaluate associated effects on the heart. We thus quantified cardiac function and central hemodynamics *in vivo* as well as SMC contractility and biaxial passive aortic properties *ex vivo*; we also used a novel computational model to associate changes in biomechanical behavior with observed microstructural features. The implications of these findings are interpreted, in part, via direct comparisons to similar data for mice that have aged normally, have germline mutations that compromise elastic fibre integrity, a feature common in central artery ageing, or have induced hypertension, which also arises in ageing.

## RESULTS

### Systolic Cardiac Function is Normal, but PWV and Diastolic Function Become Aberrant

All control (*Lmna*^*+/+*^ or *Wt*) and progeria (*Lmna*^*G609G/G609G*^ or *G609G*) mice survived to the intended 140 days (d) of age, though progeria mice were significantly smaller after ∼42d (Fig. 1A): for example, body mass at 140d was 12.1±0.8 g in female and 13.8±0.9 g in male *G609G* mice compared with 25.0±1.1 g in female and 30.4±2.2 g in male *Wt* mice (*p* < 0.01). The heart was similarly smaller in progeria, yet myocardial microstructure appeared normal (Fig. 1B). *In vivo* measurements at 140d revealed that left ventricular mass and diameter, cardiac output, and stroke volume were all significantly less in progeria (Table S1), but these metrics followed allometric scaling with body mass (*y* = *cM*^*k*^, with *M* mass and *c* and *k* allometric parameters), indicative of normal function for a smaller mouse (Fig. 1C-F). Though tail-cuff blood pressure was lower in progeria (Fig. 1G), ejection fraction and fractional shortening were similar between progeria and controls (Fig. 1H), suggesting a preserved systolic function even in the peri-morbid period (*G609G* mice died by ∼150d). In contrast, diastolic function was compromised near the end of life. *E* (peak filling velocity in early diastole) and *A* (atrial contraction induced filling velocity in late diastole) were lower in progeria mice at 140d, with *E/A* higher suggestive of diastolic dysfunction (Table S1). Moreover, *E’* was lower but *E/E’* was elevated significantly (Fig. 1I), confirming late-stage diastolic dysfunction.

**Figure 1.**
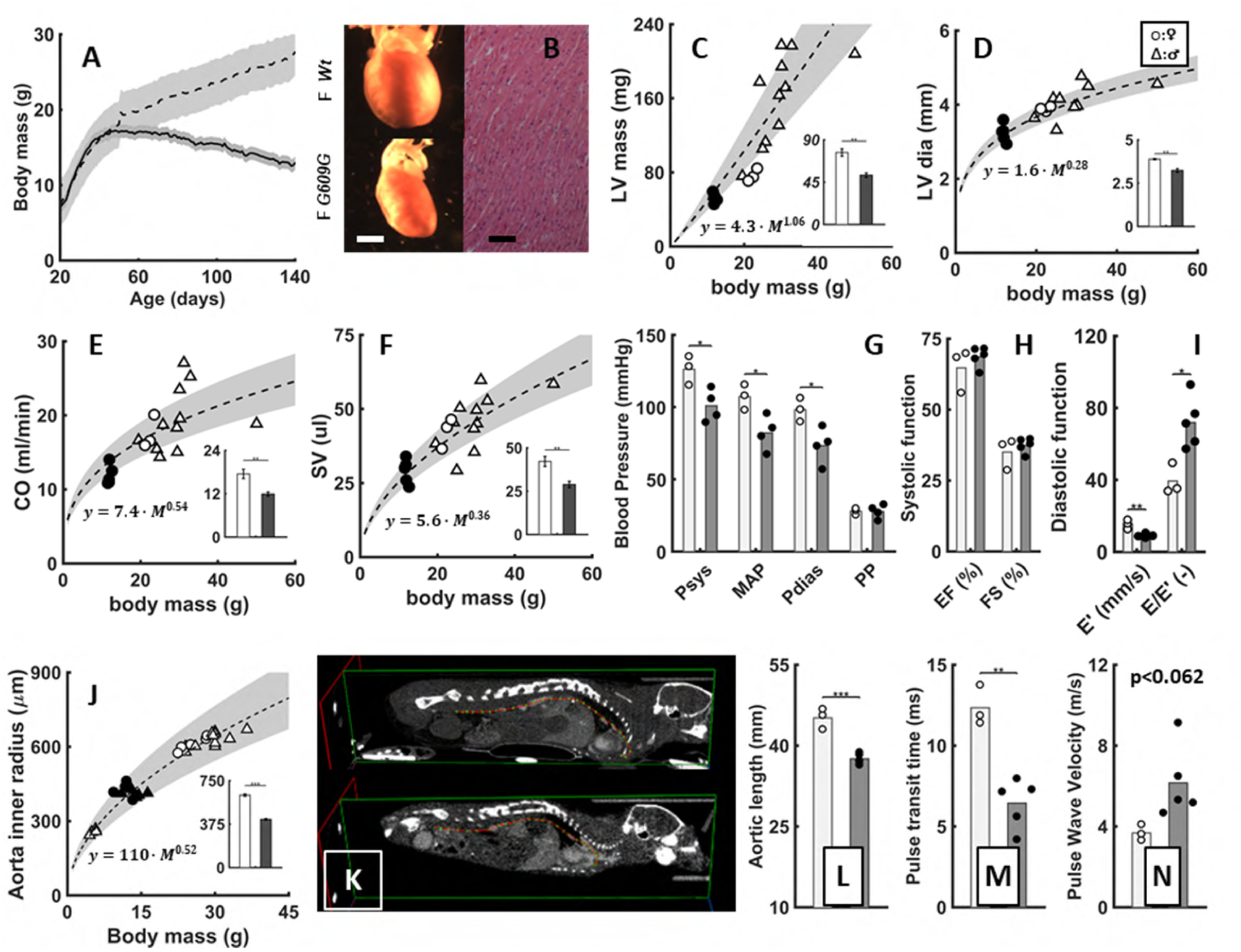
Allometric scaling reveal normal cardiac function, but PWV and diastolic Function is abnormal in progeria. **A:** Changes in body mass in female+male *Wt*=*Lmna*^*+/+*^ (dashed curve) and *G609G*=*Lmna*^G609G/G609G^ (solid curve) mice from 20 to 140 days, with 95% confidence intervals in grey. **B:** Representative images of a *Wt* and *G609G* heart at 140 days, with H&E stained cross-sections. **C-F:** Statistical differences in cardiac metrics (subfigure bar-plots) for female progeria (filled bars, circles) vs. control (open bars, circles) mice and yet appropriate allometric scaling (*y* = *cM*^*k*^, dashed lines including additional controls as open triangles; Ferruzzi et al., 2018) of (**C**) Left ventricular (LV) mass, (**D**) LV inner diameter (dia), (**E**) Cardiac output (CO), and (**F**) Stroke volume (SV). **G:** Significant reductions in systolic (Psys), mean (MAP), and diastolic (Pdias) blood pressure in 140-day-old progeria mice, but a maintained pulse pressure (PP) based on tail-cuff measurements. **H:** Ejection fraction (EF) and fractional shortening (FS) confirm LV systolic function in 140-day-old female *G609G* mice while **I:** mitral annular velocity during early filling (E’) and the ratio of early transmitral flow velocity E to E’ (E/E’) reveal LV diastolic dysfunction in the same. **J:** Loaded inner radius of the descending thoracic aorta at Psys, again compared with allometrically scaled data from additional wild-type controls. Micro-CT imaging (**K**) revealed that (**L**) the centerline distance (AL) between the aortic root and the aortic bifurcation of 140-day old female *G609G* mice was less than that in *Wt* mice. **M:** Doppler flow velocity measurements revealed that the pulse transit time (PTT) between the aortic root and aortic bifurcation was considerably less in these progeria mice, thus resulting in (**N**) an increase in pulse wave velocity calculated as AL/PTT, suggesting increased central artery stiffness. ***: P<0.001, **: P<0.01, *: P<0.05.

Proximal aortic diameters were less in progeria mice, as expected of a smaller mouse, but this metric again followed allometric scaling with body mass (Fig. 1J). Aortic lengths were less in progeria (Fig. 1K,L), but the transit time for the pulse pressure wave to travel from the aortic root to the aortic bifurcation was considerably less (Fig. 1M). Consequently, PWV was dramatically higher in late-stage progeria (Fig. 1N), trending toward significance (6.16 m/s vs. 3.68 m/s; *p* < 0.062) and suggesting increased central artery stiffness consistent with diastolic dysfunction.

### SMC Contractile Dysfunction is Progressive, Becoming Extreme

The *ex vivo* vasoactive responses were independent of sex, hence data from females and males were combined for clarity (Tables S2,S3). All four aortic segments (ascending and descending thoracic, suprarenal and infrarenal abdominal) from control mice vasoconstricted strongly to 100 mM KCl and 1 μM phenylephrine (Figs. 2A-D,S1), with 10–25% reductions in diameter under physiologically relevant, axially isometric (fixed *in vivo* axial stretch) and isobaric (90 mmHg) conditions. In contrast, vasoconstriction was attenuated at 100d and absent at 140d in all four segments in progeria. Consequently, SMC contraction reduced mean circumferential wall stress *σ*_*θ*_ (= *Pa*/*h*, with *P* pressure, *a* luminal radius, and *h* wall thickness) by 20–50% in the controls, depending on aortic segment, but not in progeria (Tables S2,S3). Hence, much-to-all of the hemodynamic loading has to be borne by extracellular matrix in late-stage progeria. Because reduced vasoconstriction could result from a loss of SMCs or compromised cellular contractility, or both, cell number was determined from histology. This quantification confirmed SMC drop-out (Figs. 2E-G). Vasoconstrictive strength was also normalized by cell number and circumference to account for different vessel calibers by age, region, and genotype, which further revealed a reduced ability of the remnant cells to contract in progeria (Fig. 2H).

**Figure 2.**
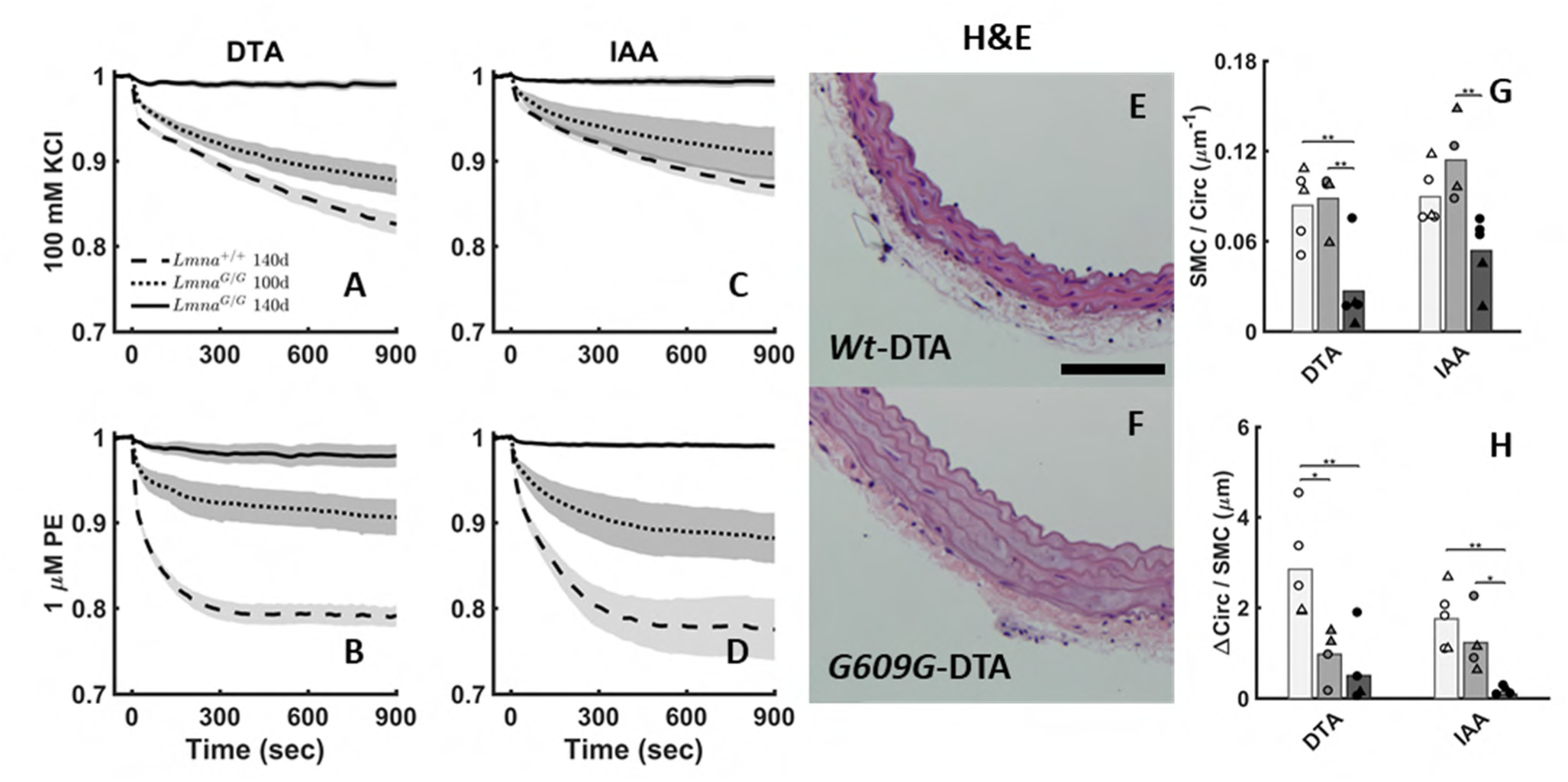
SMC contractile dysfunction is progressive, while SMC drop-out occur in late-stage. Normalized reduction in outer diameter during *ex vivo* biaxial (isobaric-isometric) vasoconstriction in response to (**A,C**) 100 mM KCl (membrane depolarization) and (**B,D**) 1 μM phenylephrine (GPCR-RhoA-ROCK pathway), at 90 mmHg and group-specific values of *in vivo* axial stretch of the descending thoracic aorta (DTA) and infrarenal abdominal aorta (IAA) from 140-day old mixed-sex *Wt* (dashed lines), and 100- (dotted lines) and 140- (solid lines) day old *G609G* mice. **E,F:** Representative histological images of 140-day *Wt* (**E**) and *G609G* (**F**) DTA stained with H&E; IAA similar. **G:** Cross-sectional smooth muscle cell (SMC) nuclei count normalized by the loaded (90 mmHg and *in vivo* axial stretch) inner circumference for the DTA and IAA for 140-day old *Wt* (open) and 100- (grey) and 140- (black) day old female (o) and male (Δ) *G609G* mice. **H:** Change in circumference (distance) per SMC, calculated as ΔCirc/SMC = Δ*d*_*o,act*_/*d*_*o*_· 2*πR*_*i*_ *λ*_*φ*_/*N*_SMC_ where Δ*d*_o,act_/*d*_o_ is the normalized reduction in outer diameter in response to 1 μM PE after 900 seconds of contraction, 2*πR*_*i*_ *λ*_*φ*_ is the loaded inner circumference, and *N*_SMC_ is the SMC nuclei count. ***: P<0.001, **: P<0.01, *: P<0.05.

### The Aortic Biomechanical Phenotype is Progressive, Becoming Extreme

Geometrical and passive mechanical data were quantified for progeria and age-matched control aortas, with data from males and females again similar (Fig. S2). Biaxial structural stiffening manifested in all four aortic regions in progeria at 100d, which progressed to extreme at 140d (Figs. 3A-D) consistent with the measured increase in PWV (Fig. 1N). Albeit not shown, all biaxial data, progeria and control, were fit well by the same nonlinear constitutive relation (Methods); associated best-fit values of the constitutive parameters are in Tables S4,S6. Combined with appropriate geometrical data (Fig. 3I,J, Tables S5,S7), these results allow one to compute key metrics such as mean biaxial wall stress (Figs. S3,S4), wall stretch (Fig. 3E,F), and material stiffness (Fig. 3G,H), and similarly the capacity of the aortic wall to store elastic energy upon deformation (Fig. 3K). Indeed, one can compute segmental PWV (Fig. 3L,S2T), which revealed significant elevations in proximal and especially distal regions. Whereas decreased circumferential stretch and wall stress were expected because of the marked thickening of the wall in progeria (Fig. 3J), the ubiquitous decrease in circumferential material stiffness and the progressive decrease in elastic energy storage were extreme (Fig. 3G,K). In the ascending aorta, for example, circumferential stiffness fell from ∼2.54 MPa in controls to 0.65 MPa in progeria at 140d, and energy storage fell from ∼88.5 kPa to 8.5 kPa at 140d (Tables S5,S7). The former reveals that the increased structural stiffness, which increases PWV, results from thickening (Fig. 3J) of the wall despite a paradoxical lower intrinsic material stiffening (Fig. 3G); the latter reveals a marked loss of mechanical functionality in late-stage progeria since a primary mechanical function of the aorta is to store elastic energy during systole and to use this energy during diastole to augment blood flow.

**Figure 3.**
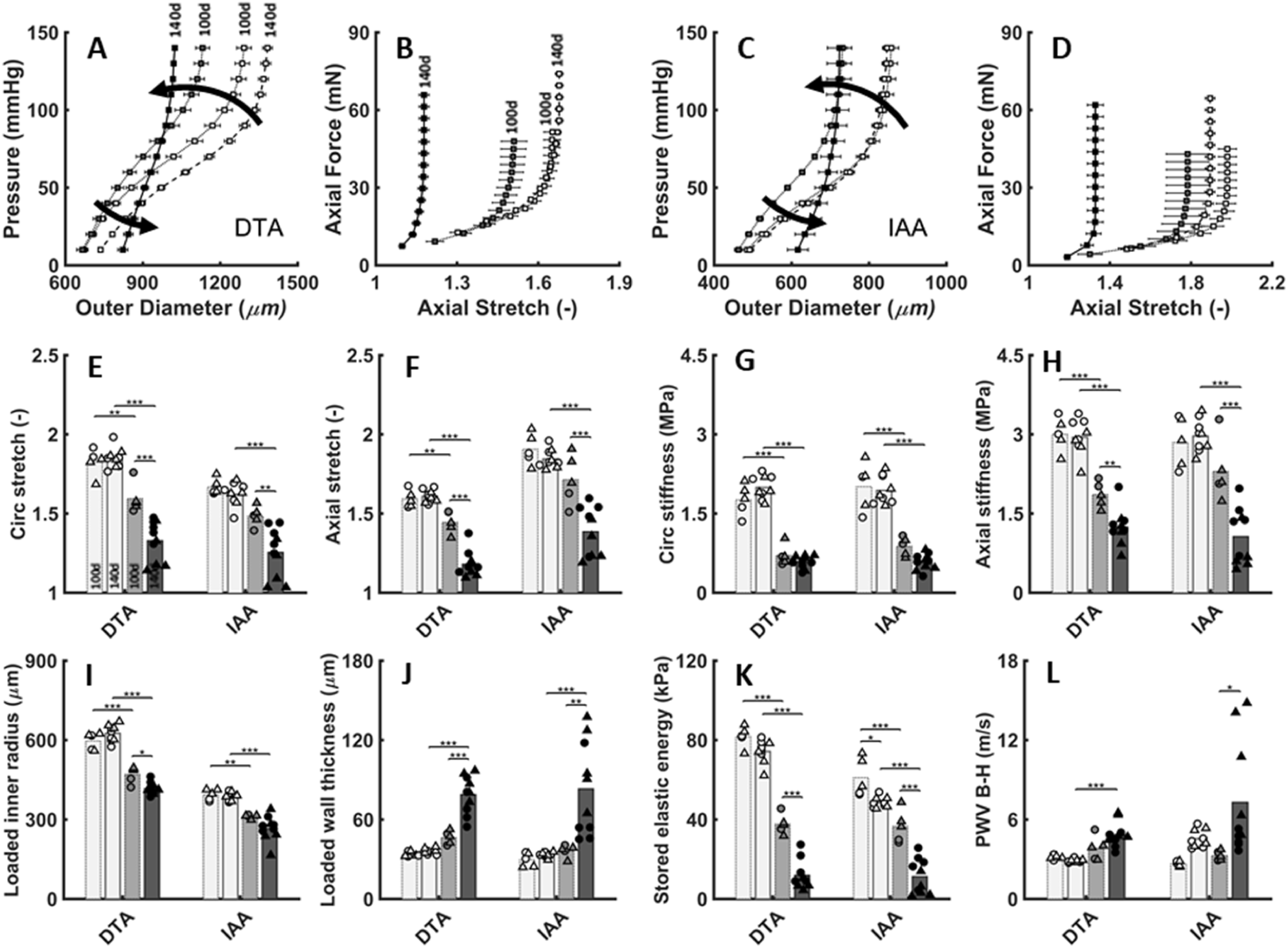
Progression of aortic biomechanical phenotype in progeria. Pressure–Outer diameter (distensibility) and Axial force–Stretch (extensibility) relationships for 100- (dotted, open) and 140- (dashed, open) day-old mixed sex (□) *Wt* as well as 100- (dotted, grey) and 140- (solid, black) day-old mixed sex *G609G* mice: **A,B -** descending thoracic aorta (DTA) and **C,D -** infrarenal abdominal aorta (IAA). **E-L:** Computed geometrical and mechanical metrics from the passive tests on the 100- and 140-day-old female (o) and male (Δ) control (open) and progeria (filled) aortas under *in vivo* systolic conditions. **E-F:** Biaxial stretch and (**G,H**) biaxial stiffness. Similarly, (**I,J)** loaded inner radius and wall thickness, (**K)** stored elastic energy *W* and (**L**) Bramwell-Hill pulse wave velocity 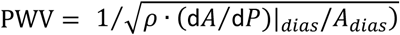, where *ρ* is the fluid mass density, *A* the lumenal cross-section, and *P* the pressure. Note that structural stiffening is revealed by the marked left-ward shifts within the physiological ranges of pressure-diameter and axial force-stretch responses, with associated losses in distensibility (circumferential) and extensibility (axial) reducing energy storage. See Figs. S2-S4 for additional regions and metrics. ***: P<0.001, **: P<0.01, *: P<0.05.

### Late-Stage Proteoglycan Accumulation is Diffuse and Dramatic

Aortic properties and function stem from the underlying microstructural composition and architecture. Despite the progressive loss of elastic energy storage capability, the elastic laminae appeared intact throughout the aorta in progeria, though more separated and less undulated at 140d (Figs. 4A-D,Q). The latter appeared to be caused by a dramatic increase in proteoglycans in progeria, especially in the media (Figs. 4A-D,M,N,S5). Immunostaining revealed that, although absent in the controls, aggrecan was surprisingly the primary proteoglycan within the media in progeria at 140d (Figs. 4E-H,O). It appears that the strongly negative fixed charge density of the proteoglycans and associated Gibbs-Donnan swelling increased the separation of the elastic lamellae and decreased their undulation. Although less dramatic, proteoglycans similarly appeared to thicken the adventitia (Fig. 4N,P) and decrease the collagen fibre undulation (Figs. 4R,S5). Finally, plotting mechanical metrics versus proteoglycan fraction revealed that the dramatic decrease in circumferential material stiffness occurred before proteoglycan production (Figs. 4S,S6), whereas the degree of change from compromised to extreme in most other metrics correlated with proteoglycan presence, which saturated at contents above ∼15%.

**Figure 4.**
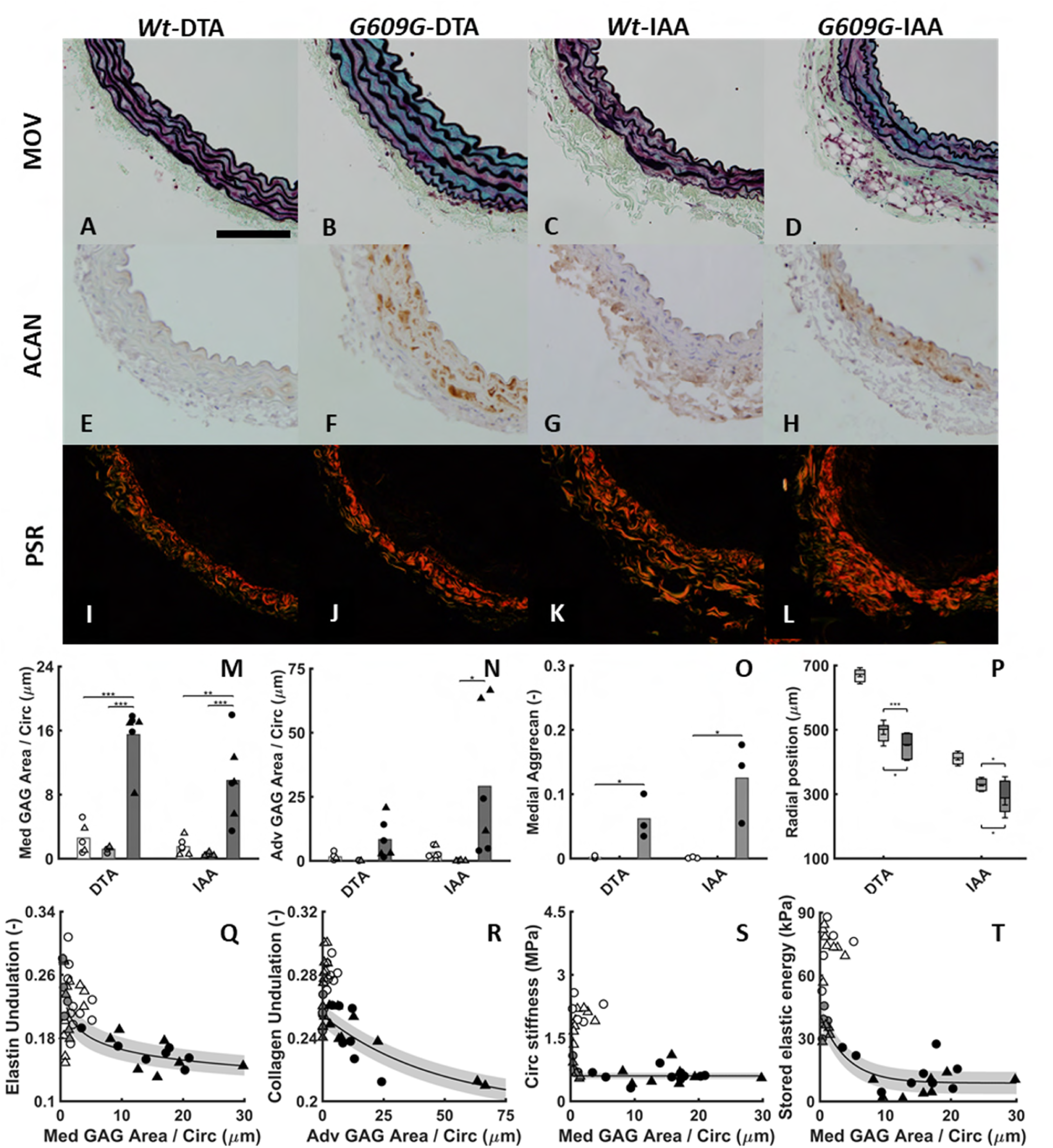
Dramatic late-stage proteoglycan accumulation in progeria aorta. Histological images of the DTA (**A,B,E,F,I,J**) and IAA (**C,D,G,H,K,L**) from 140d *Wt* (**A,E,I,C,G,K**) and *G609G* (**B,F,J,D,H,L**) mice: Movat (**A-D**), Aggrecan (**E-H**), and Picro-Sirius Red (**I-L**). Quantification of (**M**) medial GAG and (**N**) adventitial GAG, each normalized by inner vessel circumference at systolic loading conditions to account for different aortic size by region, sex, size, age, and genotype, and (**O**) fraction of medial aggrecan. See also Fig. S5. (**P**) Changes in medial (lower rectangle) and adventitial (upper rectangle) thickness driven by GAG accumulation help maintain an allometrically functional luminal radius at systolic loading conditions. See also Fig. S6 for loaded inner radius as a function of medial GAG. (**Q**) Elastin and (**R**) collagen fibre undulation (using CT-FIRE) shown as a function of medial and adventitial GAG, respectively, again normalized by loaded inner circumference at systole. (**S**) circumferential stiffness and (**T**) stored elastic energy as a function of medial GAG normalized by inner circumference. The curves are fit only to *G609G* data. GAG = glycosaminoglycan. ***: P<0.001, **: P<0.01, *: P<0.05.

### Computational Modelling Reveals Biomechanical Roles of Accumulated Proteoglycans and Remodeled Collagen

Biological assays such as western blots and (immuno)histochemistry provide critical information on tissue composition and some information on tissue organization, but they cannot reveal functional consequences, which are important clinically. Thus, we used a novel particle-based biomechanical model of the aorta (Methods, Table S8) that allows one to delineate constituent-specific consequences of compositional and organizational changes (Ahmadzadeh et al., 2018). Fig. 5A shows a histologically defined cross-section of the unloaded descending thoracic aorta from a progeria mouse, while Fig. 5B shows a general computational model of the same. The control model (not shown) was endowed with properties consistent with health; the 140d progeria model included a modified medial layer that contained five intact, nearly straight elastic laminae with dysfunctional intra-lamellar SMCs (no contractility), remodeled fibrillar collagen, and excessive stochastically distributed proteoglycans plus an adventitial layer that consisted primarily of remodeled fibrillar collagen with little proteoglycan (Fig. S7). Individual “particles” within each computational model allowed fine spatial resolution (∼2 μm in the pressurized state) and enabled the different constituents to interact physically.

**Figure 5.**
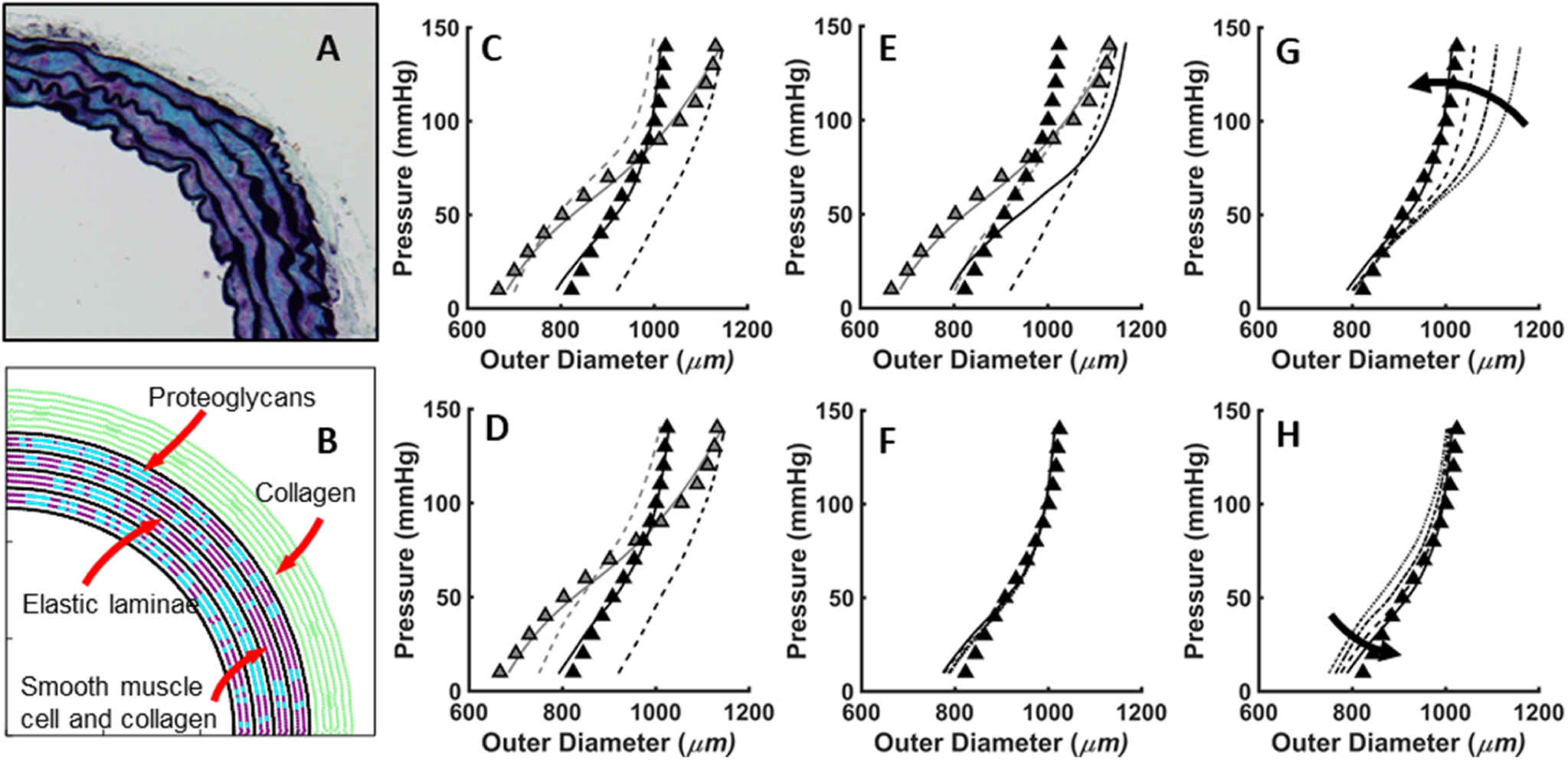
Computational modelling reveals biomechanical roles of accumulated proteoglycans and remodeled collagen. **A:** Movat-stained descending thoracic aorta (DTA) from a 140-day-old progeria mouse. **B:** Particle-based computational model of the same with stochastic distributions of proteoglycan (blue) between the elastic laminae (black) in the media, with separate mechanical properties prescribed for the primary structural constituents in both the media and adventitia, including Gibbs-Donnan swelling of proteoglycans. (**C,D**) Computational results show a good fit to the pressure-diameter response of a 100-day-old DTA (grey Δ), as well as the ability of the model to fit data for the 140-day-old DTA (black Δ) only if combining three factors (black and grey solid curves): the measured decrease in axial stretch (dark dashed), GAGs occupying 30% of the media, and remodeled collagen (light dashed) by changing either collagen (**C**) undulation or (**D**) material properties, but not just (**E**) simply adding more collagen having normal properties to thicken the wall (solid curve) to the measured level. Finally, (**F**) shows that the particular stochastic distribution of the proteoglycans is not critical, whereas the (**G**) degree of change in the collagen remodeling (e.g., 0, 5, 10, the 15% increase in deposition stretch) or the (**H**) addition of more swollen proteoglycans (0, 10, 20, 30% increases shown) is essential for fitting the experimental data. The importance of collagen in the altered properties was also demonstrated enzymatically by del Campo et al. (2019), though not in connection with proteoglycans.

Fig. 5C,D shows the ability of this model (lines) to describe pressure-diameter data (filled Δ) at group-specific fixed axial stretches for both 100d (more compliant; grey, dashed lines) and 140d (stiffer; black, solid lines) progeria aorta. Consistent with histological assessments, the model fit the control and different progeria data assuming the same amount and properties of the elastic lamellae, noting that straightening elastic laminae has little effect on stiffness because of its nearly linear stress-stretch behavior (cf. del Campo et al., 2019). Importantly, the model could only fit the 140d progeria data when introducing three modifications relative to the 100d progeria data: a marked reduction in the *in vivo* axial stretch (from 1.4 to 1.17), which was measured directly (Tables S5, S7), a marked increase (from ∼0 to 32% of the media) in highly negatively charged medial proteoglycans (Fig. 4O), and marked remodelling of adventitial collagen, consistent with either the observed loss of undulation (Fig. S5) or change in stiffness (Fig. 5D). Simulations with reduced axial stretch alone and similarly remodeled collagen alone showed that these effects were not sufficient individually; rather all three changes needed to co-exist to capture the structurally stiffer behavior (solid line). Similarly, simply increasing the thickness of the collagen-rich adventitia with similar collagen could not describe the 140d data (Fig. 5E). Finally, Fig. 5F shows that it was not the specific (stochastic) distribution of the proteoglycans that mattered, but rather the associated degree of collagen remodelling (Fig. 5G) plus the proteoglycan accumulation (Fig. 5H) at 140d.

### Comparisons Against Normal Ageing Highlight the Late-Stage Severity of Progeria

Fig. 6 compares key biomechanical metrics at group-specific systolic pressures for the descending thoracic aorta of male mice across six different models (see also Fig. S8). Note the consistency across controls for different ages of normal young adults, namely 56-, 84-, and 140-day-old C57BL/6 mice, particularly compared with 350- and 700-day-old mice that aged naturally (Ferruzzi et al., 2019) and the 100- and 140-day-old progeria (*Lmna*^*G609G/G609G*^) mice (Fig. 6A-D). The 140d progeria mice exhibited the most dramatic phenotype across all metrics, but interestingly the 100d progeria mice shared many similarities with the 700-day old naturally aged mice, especially wall thickness, distensibility, extensibility, and energy storage, though not material stiffness, which was lower in progeria. Albeit not shown, the control groups at 56 to 140 days of age were very similar to 140-day-old littermate controls (*Lmna*^*+/+*^), which were used for comparison in Fig. 6E-H. Thus consider direct comparisons of aortic data for 700-day-old naturally aged (repeated) mice as well as 56- and 90-day-old Marfan syndrome (*Fbn1*^*mgR/mgR*^) mice (Bellini et al., 2017), 140-day-old fibulin-5 deficient (*Fbln5*^*-/-*^) mice (Ferruzzi et al., 2015), and 84-day-old angiotensin II infusion induced hypertensive C57BL/6 mice (Bersi et al., 2016) – again versus 100d and 140d progeria data. The 140d progeria aortas again exhibited the most severe phenotype of all models, particularly with dramatic reductions in biaxial wall stretch (distension and extension), wall stress, and material stiffness. Only the reduced elastic energy storage in the induced hypertension model approached that of the 140d progeria aorta, noting that these hypertensive aortas thickened dramatically due to adventitial fibrosis and thus resulted from very different microstructural changes (cf. Fig. 4). Albeit not shown, the contractile response to high KCl or phenylephrine was also lowest in the 140d progeria aorta of all of the models considered here (*Wt*, fibulin-5 null, and induced hypertension), consistent with a previously reported low contractile protein expression in progeria (Capell et al., 2008).

**Figure 6.**
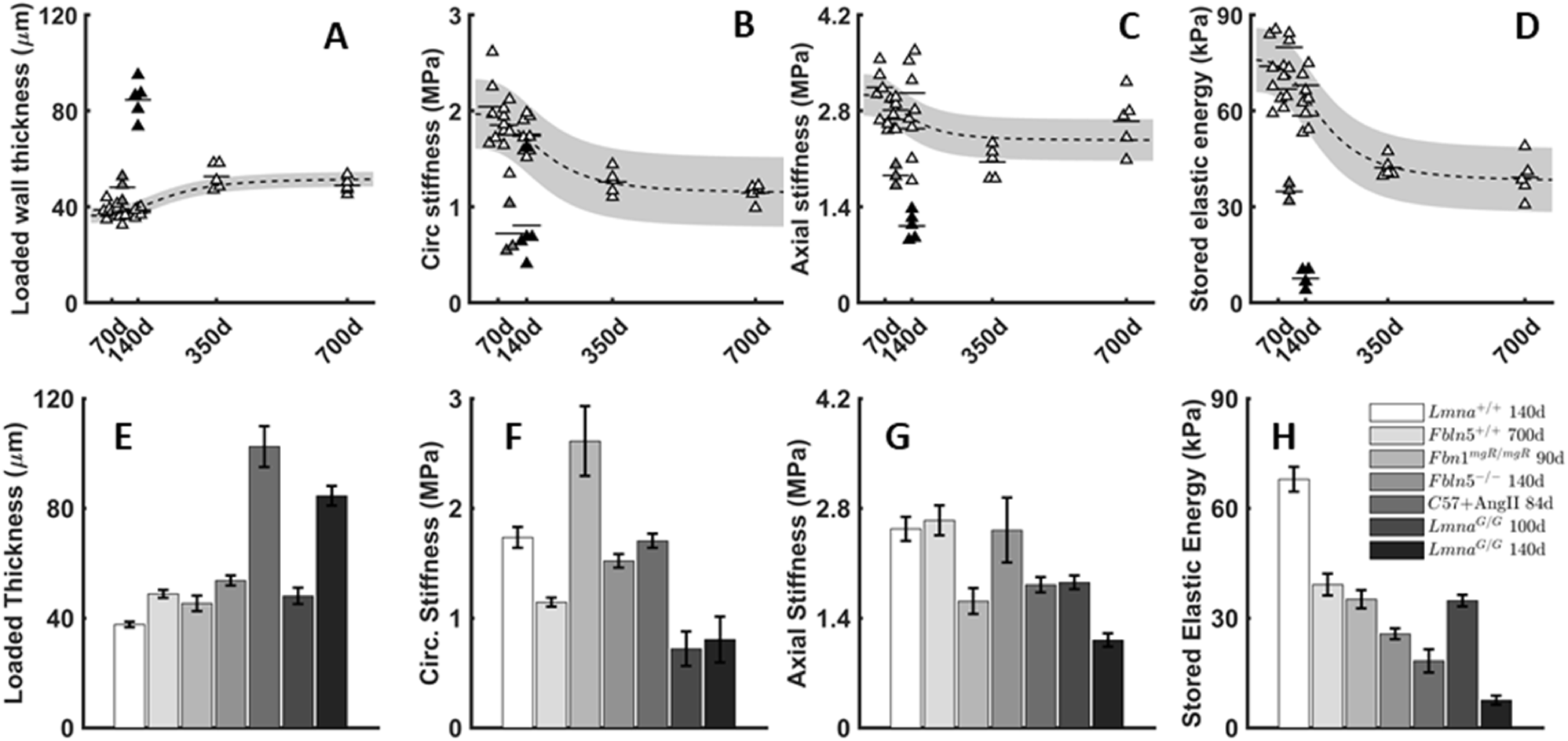
Comparison of vascular metrics in progeria and normal ageing. Key geometrical and mechanical metrics for the descending thoracic aorta from male mice as a function of (**A-D**) natural aging and (**E-H**) aging-related disease conditions, each calculated at group-specific systolic pressures and *in vivo* axial stretch. (**A,E**) wall thickness, (**B,F**) circumferential material stiffness, (**C,G**) axial material stiffness, and (**D,H**) elastically stored energy; see Supplemental Fig. S8 for additional metrics. In A-D, data for male wild-type controls at 56d, 84d, 140d, 350d, and 700d of age (open Δ, grey confidence region) are contrasted with those from 100- (grey Δ) and 140- (black Δ) day old *G609G* mice. In E-H, data from mice naturally aged to 700d (repeated from A-D) as well as those that are fibrillin-1 deficient (*Fbln1*^*mgR/mgR*^), fibulin-5 null (*Fbln5*^*-/-*^), or hypertensive (C57+AngII) are again compared with 100d and 140d old *G609G* mice; severe elastopathies and hypertension are both common in aortic aging. As it can be seen, the 100-day old *G609G* mice exhibit an aortic phenotype that is similar in different ways to those in extreme normal aging (to 700 days) or with severely compromised elastic fiber integrity (*Fbln5*^*-/-*^) or hypertension, yet the 140-day old peri-morbid *G609G* mice have the most severe aortic phenotype of all mice.

## DISCUSSION

HGPS is an ultra-rare disorder, presenting in 1 out of 4–8 million live births. Because patient data are scant, mouse models provide critical insight into the pathogenesis of this devastating syndrome as well as into mechanisms and manifestations of vascular ageing in general. There are numerous reports of central artery ageing in *Wt* mice. A comparison of 6-versus 20-month-old C57BL/6 mice revealed 1503 differentially expressed genes in the thoracic aorta, with significant differences in calcium handling, extracellular matrix, and cell adhesion as well as increased (23%) aortic PWV with consequent increased (17%) central blood pressure (Rammos et al., 2014). Structural stiffening of normally aged murine arteries has been ascribed largely to adventitial fibrosis (Fleenor et al., 2010), and adventitial thickening manifests in the c.1824C>T;p.G608G transgenic mouse model of progeria (Varga et al., 2006) and in some (Kim et al., 2018) though not all (del Campo et al., 2019) central arteries in *Lmna*^*G609G/G609G*^ mice. Although clinical phenotypes are often associated with gene expression or histological findings, one must also quantify associated functional metrics that define the aortic phenotype, which ultimately governs cardiovascular health and disease risk (Laurent and Boutouyrie, 2015; Humphrey et al., 2016). Indeed, although it has long been speculated that mechanical stress plays a key role in the disease (Varga et al., 2006; Olive et al., 2010; Kim et al., 2018), there had not yet been any detailed biomechanical analysis.

It was instructive to compare the present biomechanical results directly with those for normal ageing (Ferruzzi et al., 2018) as well as with those for compromised elastic fibre integrity (Bellini et al., 2017; Ferruzzi et al., 2015), a feature common in aortic ageing, and those for hypertension (Bersi et al., 2016; Korneva and Humphrey, 2019), which often associates with ageing. Remarkably, the progeria phenotype at 100d was similar to that of both elastic fibre compromised- and naturally aged- (to 700 days) aortas, indicative of highly accelerated ageing in progeria. Nonetheless, there was a dramatic worsening of the cardiovascular state from 100d to 140d in progeria, consistent with the histological appearance of extensive proteoglycans, resulting in an aortic biomechanical phenotype that was extreme relative to natural ageing, severe elastopathies, and hypertension, each of which also increase PWV and compromise diastolic heart function. Although aortic thickening was most dramatic in angiotensin II–induced hypertension, the associated reduction in wall stress *σ*_*θ*_ was greatest (∼75%) in the 140d progeria aorta (Fig. 3) due to the lower blood pressure, smaller lumen, and markedly thickened wall (with *σ*_*θ*_ = *Pa*/*h*). Reduced peripheral blood pressure excluded hypertension as a driver of the phenotype. Importantly, allometric scaling (cf. Korneva et al., 2019) revealed that the smaller lumen (Fig. 1J) was appropriate for smaller mice having reduced cardiac output (Fig. 1E), hence excluding pathologic inward remodelling in progeria though suggested elsewhere (del Campo et al., 2019). The greater reduction in axial (pre)stretch in progeria (∼28%; Table S5) also contributed to these differences, emphasizing the need for biaxial measurements, which had not been performed previously. Axial pre-stretch normally arises in arteries due to perinatal deposition and cross-linking of elastic fibres having a long half-life, but increases in other matrix constituents can reduce its extent in maturity. Notably, dramatic increases in intramural proteoglycans late in progeria contributed to both wall thickening and the reduced axial recoil, which computational modelling revealed as fundamental to the marked alterations in wall mechanics (Fig. 5).

We previously showed that decreases in elastic energy storage upon pressurization reflect a loss of mechanical functionality of the aorta (Ferruzzi et al., 2015), which should store energy during systole and use this energy during diastole to work on the blood (Wagenseil and Mecham, 2009). Such decreases are common in cases of compromised elastic fibre integrity and hypertension (Bellini et al., 2017; Korneva and Humphrey, 2019), though for different reasons – an inability to store energy in impaired elastic fibres in the former and an inability to distend competent elastic fibres in the latter. Remarkably, despite preserved elastic laminae (Figs. 4,S5), the 140d progeria aorta exhibited the greatest loss of energy storage capability of all models studied. The associated reduced distensibility of the structurally stiffer wall, and thus inability to deform the elastic fibres, again appeared to result primarily from the excessive intramural proteoglycan at 140d (Figs. 4,5). Critically, the extreme loss of energy storage by the ascending aorta (Fig. S2) would compromise diastolic cardiac function further, for energy stored in this segment can help lift the base of the heart and promote diastolic filling (Ferruzzi et al., 2018). Whereas increased circumferential material stiffness correlates with thoracic aortic aneurysm, as in Marfan syndrome (Bellini et al., 2017; Korneva et al., 2019), perhaps most notable herein was its dramatic reduction in the 100d and 140d progeria aortas. An inability to control circumferential stiffness appears to reflect a compromised ability of the intramural cells to mechano-sense and mechano-regulate their local mechanical environment (Humphrey et al., 2015), consistent with observed changes in matrix and adhesion molecule gene expression in natural ageing (Rammos et al., 2014). Importantly, the markedly lower intrinsic aortic stiffness in progeria is also consistent with the finding that normal levels of lamin A scale with tissue stiffness (Swift et al., 2013). Whereas prior reports show that decreased stiffness of the surrounding matrix can drive lower lamin A in normal cells, our findings suggest further that loss of normal lamin A or presence of progerin can compromise mechanical homeostasis and manifest as a lower matrix stiffness. It is thus critical to delineate intrinsic material stiffness (Fig. 3G) and overall structural stiffness (cf. del Campo et al., 2019), which results from a combination of material stiffness and wall thickness and ultimately affects the hemodynamics (Fig. 3L). That progeria results in a paradoxically reduced material stiffness but greater structural stiffness is consistent with both the *Lmna* mutation and clinical phenotype.

Recalling that progeria patients exhibit increased PWV (Gerhard-Herman et al., 2012), the diffusely increased local structural aortic stiffness in the 140d progeria aortas (Fig. 3L) manifested globally as an increased PWV (6.16 m/s late in progeria), which was greater than in severe elastopathy (4.47 m/s in fibulin-5 deficiency; Cuomo et al., 2019) and normal murine ageing (3.76 m/s; Rammos et al., 2014), all relative to control (3.04 m/s). That PWV increased despite the markedly reduced circumferential material stiffness highlights the important structural effect of wall thickening, particularly when combined with an allometrically reduced luminal caliber 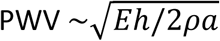, with *E* a tensile material stiffness, *h* wall thickness, *a* luminal radius, and *ρ* blood viscosity; Cuomo et al., 2019). That computed values of PWV differed slightly by aortic region may explain why the globally measured value did not reach significance (Fig. 1N) while highlighting the importance of local mechanobiology on global physiology (Humphrey et al., 2016), which in this case manifested as a compromised diastolic cardiac function (Fig. 1).

It was somewhat remarkable that systolic function was largely preserved, even in the peri-morbid period (140d), consistent with the original report at a younger (103d) age (Osorio et al., 2011). We emphasize two important points here. First, because of the significantly smaller body mass *M* in progeria after about 42d (Fig. 1A), allometric scaling (*y* = *cM*^*k*^) is essential when assessing non-scaled cardiovascular metrics (Korneva et al., 2019), including cardiac output, stroke volume, and aortic dimensions, each of which were statistically lower in progeria but otherwise appropriate for the smaller mice. Interestingly, our best-fit allometric parameters *k* were comparable to those in other rodents (White et al., 1968) as well as humans (de Simone et al., 1997; Dewey et al., 2008). We submit, therefore, that allometric (or similar) scaling be used when interpreting cardiovascular data from progeria patients. Second, recalling that diastolic dysfunction manifests in progeria patients (Prakash et al., 2018), the impaired diastolic function in the 140d progeria mice (trend toward higher *E/A* and statistically higher *E/E’*) may have been underestimated due to the anesthesia during echocardiography. Such cardiac deficits could be exacerbated by physical exertion, perhaps explaining the progressively sedentary lifestyle of these mice.

We quantified absolute histological changes but normalized by luminal circumference to account for differences in caliber due to age, sex, body size, genotype, and location along the aorta. Considered in this way, the density of elastic laminae dropped from 100d to 140d in progeria, consistent with increases in collagen and especially proteoglycans, but remained higher than in *Wt* (Fig. S6A), likely due to the allometrically smaller lumen. Thus, loss of energy storage capability was not due to compromised elastin. These as well as the computational findings suggest further that the adverse aortic remodelling began after the developmental period when elastin is deposited (Wagenseil and Mecham, 2009). Given the lower blood pressures and thus lower wall stress and stiffness during development, it seems that the SMCs fashioned appropriate elastic fibre integrity, consistent with the lack of any phenotype in compliant, low-stressed organs in progeria (e.g., brain and liver; Kim et al., 2018). In contrast, collagen turns over continuously (Valentin and Humphrey, 2009). Collagen density remained nearly normal in progeria (Fig. S5), yet turnover of developmentally normal collagen in the face of higher blood pressures in maturity explains some of the progressive worsening of the aortic phenotype, recalling that the computational model predicted that the observed collagen differed mechanically in late-stage progeria. Although the reduced undulation of adventitial fibres (Figs. 4,S5) might have been expected to increase material stiffness, the thickened media likely stress shielded these fibres at normal pressures (Bellini et al., 2014), though they could yet help prevent overall dilatation, noting that aneurysms have not been reported in progeria. That the proteoglycans thickened the media, separating the elastic lamellae via Gibbs-Donnan swelling, could have compromised SMC mechano-sensing further (Roccabianca et al., 2014), thus adversely affecting matrix remodeling and cell adhesion (cf. Rammos et al., 2014) and perhaps inducing cell death via anoikis (Meredith et al., 1993). Associated consequences could be greater at higher blood pressures, which would decrease exercise tolerance consistent with the sedentary life-style of the older progeria mice.

When appropriately normalized, histology also suggested that the lost vasoconstriction in the peri-morbid period (Fig. 2A-D) resulted both from the previously reported SMC drop-out (Villa-Bellosta et al., 2013) and the new finding that the contractile capacity was lower per remnant cell, mainly for phenylephrine, an alpha1-adrenergic agonist. Phenylephrine mediates contraction through the RhoA-ROCK pathway, consistent with reports of reduced RhoA activity in HGPS mice and the hypothesis that a disrupted LINC complex correlates with weakened cell-matrix adhesion (Hale et al., 2008). Reduced central artery contractility manifests in many conditions, including Marfan syndrome (Eberth et al., 2009), fibulin-5 deficiency (Murtada et al., 2016), induced hypertension (Korneva and Humphrey, 2019), and natural aortic ageing (Wheeler et al., 2015). Yet, vascular contractility is only attenuated, not absent, in these other cases. Varga et al. (2006) previously reported diminished *in vivo* vasodilation to sodium nitroprusside in the transgenic c.1824C>T;pG608G mouse, while Capell et al. (2008) reported a decrease in SM-α actin, yet ours is the first report of a total, late-stage loss of vasoconstrictive capacity under physiological conditions. Reduced vessel-level contractility compromises the ability of the cells to maintain or remodel the wall in response to hemodynamic loads (Dajnowiec and Langille, 2007; Valentin et al., 2009). Reduced vasoconstriction also suggests a loss of cell-tissue level actomyosin activity, which is needed to mechano-sense and mechano-regulate the extracellular matrix (Humphrey et al., 2015). It should not escape one’s notice, therefore, that the altered nuclear stiffness, severely diminished actomyosin activity, and compromised extracellular matrix in progeria lie along the mechanotransduction axis, suggesting that progressive microstructural changes follow a worsening ability of the SMCs to remodel or repair the matrix, resulting in excessive proteoglycan production, particularly aggrecan which is typically not found in the aorta (cf. Cikach et al., 2018) though it is produced by dermal fibroblasts in HGPS (Lemire et al., 2006).

Increased PWV, reflective of central artery stiffening, has emerged as characteristic of the systemic vasculature in progeria (Gordon et al., 2012; Gerhard-Herman et al., 2012; Prakash et al., 2018). Such stiffening in the general population is strongly predictive of future cardiovascular events, including myocardial infarction, stroke, and heart failure (Vlachopoulos et al., 2010; Mitchell et al., 2010), and may be an indicator of premature vascular ageing (Nilsson et al., 2013). Our findings revealed highly accelerated aortic aging in progeria up to 100 days, comparable to that of natural aging and genetically compromised elastic fibre integrity, followed by an extreme late-stage (100-to-140d) loss of aortic contractile and biomechanical function, independent of sex, leading to significant increases in structural stiffness and PWV despite a paradoxically low intrinsic material stiffness consistent with the lamin A mutation and indicative of compromised mechanical homeostasis. Because the aortic phenotype and associated cardiac performance are much less severe at 100d (Osorio et al., 2011) than in peri-morbid 140d progeria mice (Figs. 1-3), there may be a late therapeutic window that could expand treatment options to drugs that otherwise would be questioned due to growth restriction in children (cf. DuBose et al., 2018). In particular, it would be prudent to control SMC phenotype and limit proteoglycan accumulation, which appear to drive the extreme central artery stiffening and devastating cardiovascular sequelae in HGPS.

## EXPERIMENTAL PROCEDURES

### Animal Studies

Mice were generated using heterozygous (*Lmna*^*G609G/+*^) breeding pairs. A total of 37 female (F) and male (M) control (*Lmna*^*+/+*^ or simply *Wt*) and progeria (*Lmna*^*G609G/G609G*^ or simply *G609G*) offspring were studied at either 100 days of age (∼14 weeks) or 140 days of age (20 weeks), the latter to study the peri-morbid condition since these progeria mice died around 150 days of age. As we reported at the September 20-24, 2018 Progeria Research Foundation meeting, this extension in lifespan beyond the expected 103 days (cf. Osorio et al., 2011) was achieved by providing chow on the floor of the cages to facilitate eating as the mice became progressively weaker. We did not use a high-fat diet to extend lifespan further, however (cf. Kreienkamp et al., 2018). Conscious blood pressures were measured using a standard tail-cuff device (Kent Scientific). At the appropriate endpoint, at times following the collection of in vivo data, an intraperitoneal injection of Beuthanasia-D was used to euthanize the mice and the aorta was excised from its root to the aortic bifurcation. These excised vessels were used for ex vivo biomechanical evaluation and subsequent (immuno)histological examination as described below.

### Ultrasound and microCT

Following isoflurane anesthesia, echocardiographic and vascular ultrasonic data were collected in vivo on a sub-group of 140-day old female littermate (*n* = 2) and progeria (*n* = 5) mice, followed by microCT imageing. These data were contrasted with prior data on 20-week old wild-type female mice (*n* = 8). Noninvasive echocardiographic and vascular ultrasonic data were collected using a Visualsonic Vevo 2100 system with a linear array probe (22-55 MHz). Systolic and diastolic function of the left ventricle (LV) was quantified using standard methods (Ferruzzi et al., 2018). Briefly, B-Mode imaging in parasternal long axis (LAX) and short axis (SAX) views and M-Mode imaging in both planes tracked cavity diameter at the level of the papillary muscles to measure LV systolic function. B-Mode images of the LV outflow tract diameter and pulsed-wave Doppler images of blood velocity patterns across the aortic valve estimated aortic valve area to check for valve stenosis. LV diastolic function was monitored using an apical four-chamber view, which was achieved by aligning the ultrasound beam with the cardiac long axis. Color Doppler imaging located the mitral valve, while pulsed-wave Doppler measured mitral inflow velocity. Finally, Doppler tissue imaging from the lateral wall and interventricular septum measured velocities associated with tissue motion. B-Mode imaging also located each aortic segment via anatomical landmarks and measured ascending aortic deformations while standard M-Mode images measured time-varying luminal diameters in all four regions in both SAX and LAX views. PW Doppler measured near centerline blood velocities proximally (near the aortic root) and distally (close to the aortic bifurcation), which enabled assessment of pulse wave transit times.

The aortic path length traveled by the pulse wave (from the ascending aorta to the aortic bifurcation) was measured from an in vivo micro-CT angiogram. Briefly, following a bolus intrajugular injection of 5 ml/kg exia 160 contrast agent (Binitio Biomedical Inc), mice were imaged in a micro-CT scanner (MILabs) at 50 kV tube voltage, 0.48 mA tube current, 40 ms exposure, and 480 projections. Projections from each scan were reconstructed into 3D volumes, with a voxel size of 80×80×80 μm^3^, and arterial cross-sections along the aortic tree were segmented semi-automatically using a 2D level-set method using the open source software package *CRIMSON* (www.Crimson.org).

### Allometric Scaling

Given the markedly reduced body mass of the progeria mice at all ages beyond 6 weeks, independent of sex, we considered possible allometric scaling of the form *y* = *cM*^*k*^, where *y* is any metric of interest (e.g., cardiac output or luminal diameter), *M* is body mass (in grams), and *c* and *k* are allometric constants determined from linear regression of data from normal wild-type mice plotted as ln(*y*) = ln(*c*) + *k*ln(*M*), with ln the natural logarithm. Data from the progeria mice were then plotted with the allometric data from wild-type mice to determine congruency or not with normal scaling.

### Biomechanical Testing

We studied four aortic regions: ascending thoracic aorta (ATA), proximal descending thoracic aorta (DTA), suprarenal abdominal aorta (SAA), and infrarenal abdominal aorta (IAA). To facilitate direct comparison of the present data (*n* = 5 M and 5 F *Wt* and *n* = 5 M and 5 F *G609G* aortas at 140 days and *n* = 4 mixed-sex *Wt* and *n* = 5 mixed-sex *G609G* aortas at 100 days) with previously reported data for multiple mouse models, we used the same validated methods of biaxial testing and data analysis (Ferruzzi et al., 2015,2019). Briefly, segments from each of the four regions were isolated by blunt dissection, cannulated, and placed within our custom computer-controlled testing device within a Krebs-Ringer’s solution at 37°C (for active studies) or Hank’s buffered solution at room temperature (for passive studies). For the active tests, vessels were set at their in vivo length and pressurized at 90 mmHg (Murtada et al., 2016). The vessels were then contracted sequentially with two vasoactive agents: 100 mM potassium chloride (KCl), which depolarizes the cell membrane and increases intracellular calcium, and 1 μM phenylephrine (PE), which is an alpha adrenergic agonist. For the subsequent passive tests, vessels were preconditioned via cyclic pressurization from ∼80 to 120 mmHg while held at their individual in vivo axial length and exposed to Hank’s solution, then subjected to a series of seven biaxial protocols: cyclic pressurization from ∼10 to 140 mm Hg at three values of axial stretch (95, 100, and 105% of the in vivo value) and cyclic axial stretching at four fixed values of luminal pressure (10, 60, 100, or 140 mmHg). Data collected on-line included outer diameter, luminal pressure, axial length, and axial force.

We conclude by noting that biaxial tests, both active and passive, are critical for evaluating arterial behavior under physiologically meaningful conditions. The only prior biomechanical tests (passive) reported on vessels from progeria mice were based on wire- or pressure-myography (del Campo et al., 2019), neither of which captures the full biomechanical behavior and thus are insufficient for detailed phenotyping.

### Biomechanical Properties

Data from the unloading portions of the last cycles of the seven passive pressure-diameter and axial force-length protocols were fit simultaneously using a Marquardt-Levenberg nonlinear regression and a validated nonlinear stored energy function *W* (Ferruzzi et al., 2015; Bellini et al., 2017). This function accounts for isotropic contributions of an amorphous matrix (via a neo-Hookean function), thought to arise mainly from elastin and glycosaminoglycans (GAGs), and anisotropic contributions due to multiple families of locally parallel collagen fibres and circumferentially-oriented passive smooth muscle (via Fung-type exponential functions), namely

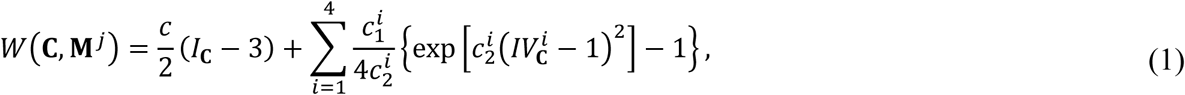

where *I*_**C**_ = *tr*(**C**) and 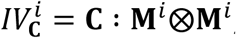, with **C** = **F**^T^**F** computed from the deformation gradient tensor **F** = diag[*λ*_r_, *λ*_0_, *λ*_z_] that is inferred directly from experimental measurements of changes in diameter and length, which define the stretch ratios *λ*_*i*_, with det(**F**) = 1 because of assumed incompressibility of the wall. The direction of the *i*^th^ family of fibres in the traction-free reference configuration is 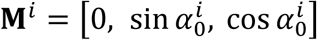, with angle 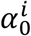 defined relative to the axial direction. Based on prior microstructural observations for wild-type elastic arteries, and the yet unquantified effects of copious cross-links amongst the multiple families of fibres, we include possible axial 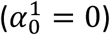, circumferential 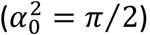, and two symmetric diagonal families of fibres 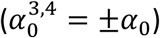. Hence, the eight model parameters are: 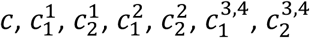, and *α*_0_. Values of biaxial stress and material stiffness are computed from appropriate differentiation of the stored energy function, then compared at physiologic pressures and individual in vivo values of axial stretch.

### Histology and Immunohistochemistry

Following ex vivo testing, samples were fixed overnight in 4% formalin while unloaded, then stored in 70% ethanol at 4°C until embedding in paraffin and sectioning at 5 µm. Sections were stained with Hematoxylin & Eosin (H&E) to count cell nuclei, Verhoff Van Gieson (VVG) to quantify elastic fibres (black), Movat’s Pentachrome (MOV) to quantify intramural proteoglycans (blue), Masson’s Trichrome to quantify cytoplasm (red) and fibrillar collagen (blue), and Picro-Sirius Red (PSR) to delineate collagen fibre size under polarized light (red to green). Slides were imaged using an Olympus BX/51 microscope, with bright- and dark-field imaging at 20x magnification. We analyzed 3 cross-sections for each sample and stain to reduce intra-specimen variability. We used previously developed custom MATLAB software to identify absolute values and area fractions of each load-bearing constituent (Bersi et al., 2016). Briefly, background subtraction and pixel-based thresholding classified each pixel within the stained sections as elastin, collagen, cytoplasm, or medial proteoglycans. Similarly, we identified the fraction of thick (red-orange) or thin (yellow-green) birefringent collagen fibres in the media and adventitia of PSR-stained sections. Medial and adventitial areas were delineated by the external elastic lamina in MOV images. Immunostaining was used to delineate the proteoglycans aggrecan and versican. Gray-scaled PSR images were analyzed with a curvelet-denoising filter followed by an automated fibre-tracking algorithm combined with (CT-FIRE) (Bredfeldt et al. 2014) to quantify undulation of collagen fibers; elastin lamellar undulation was determined from VVG-stained sections.

### Particle-based Computational Model

Whereas the aforementioned continuum model of the arterial wall enables one to compute locally averaged material properties and wall stresses while satisfying mechanical equilibrium, we also used a unique particle-based model to examine individual contributions by the different constituents, and their interactions, including elastic fibres organized into concentric laminae, single layers of smooth muscle cells, collagen fibres, and diffuse proteoglycans that exhibit Gibbs-Donnan swelling (Ahmadzadeh et al., 2018). Briefly, we defined the arterial wall as a collection of *j*=1,2…,N “particles” that are endowed with separate biophysical properties and subject to Newton’s second law of motion: 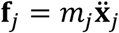, where *m* is the mass and 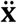 the acceleration of particle *j*. Each particle interacts directly with other particles within a locally defined neighborhood, defined by a particle list, and continuum-type quantities can be computed by introducing a smoothing (kernel) function

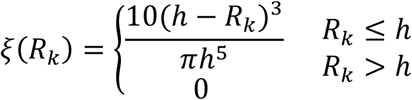

where *R*_*k*_ = |**X**_*k*_ - **X**_*j*_| denotes the distance between any particle *k* and particle *j*, and *h* defines the “kernel support”. In this way any quantity of interest *g* at original particle location **X**_*j*_ can be computed via

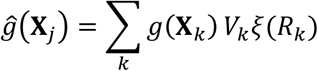

where *V*_*k*_ denotes the local material volume associated with the listed neighboring particle *k*. Further details on the theoretical framework as well as the numerical implementation on a high performance computer (Intel Xeon E5-2660 CPU running at 2.6 GHz on 128 cores) can be found elsewhere (Ahmadzadeh et al., 2018).

Briefly, by mirroring the geometry and microstructural composition of the DTA of the progeria mouse at 100 days of age at the *in vivo* homeostatic state (luminal pressure of 85 mmHg and axial stretch of 1.40), the domain of the model was a portion of cylindrical wall with inner and outer radii *R*_in_ = 434 μm and *R*_out_ = 485 μm (with wall thickness *H* = 51 μm), divided into an inner layer representing the media (with thickness *H*_M_ = 39 μm) and an outer layer representing the adventitia (*H*_A_ = 12 μm). Because the 100-day old progeria aorta has minimal proteoglycan accumulation, the wall at this stage was assumed to be devoid of added, swollen proteoglycans. The media was further divided into five alternating layers of elastin (elastic laminae), separated by intra-lamellar regions composed of smooth muscle cells and collagen. Employing our particle-based framework, this cylindrical domain was discretized into an arrangement of uniformly distributed particles (with inter-particle spacing ∼2 *μm*) that represented their surrounding space bounded by their immediate neighboring particles. The particles positioned at the elastic laminae, intra-lamellar regions, and adventitia are color coded and labeled as elastin (black) particles, intra-lamellar (red or blue) particles, and adventitial (green) particles, respectively.

Importantly, the total strain energy function at each particle *j* having contributions from any normal structural constituent (i.e., elastin, smooth muscle cells, and collagen) is

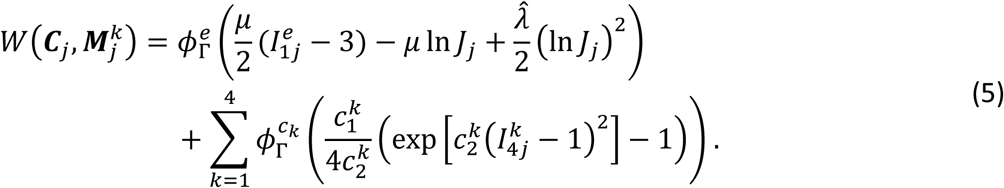

The first term represents the isotropic behavior of elastin networks, which is best described by a neo-Hookean material response with shear modulus *μ* and Lamé constant 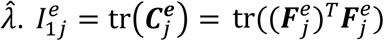 is the first invariant of the right Cauchy-Green tensor 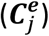 where 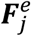 is the deformation gradient tensor for elastin (superscript *e* denotes elastin). The second term accounts for the directional behavior of collagen fibres and smooth muscle cells. The collagen fibres are categorized into four directions (or families), denoted by *k* = 1 for axial, *k* = 2 for circumferential, and *k* = 3,4 for symmetric diagonal. Smooth muscle cells are also assumed to behave in the circumferential direction, therefore, they are considered within a combined contribution with the circumferentially-oriented collagen (*k* = 2). Accordingly, 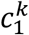 and 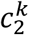 are the material parameters and 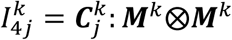 is the square of the stretch in the *k*^*th*^ family with ***M***^*k*^ = [0, sin *α*, cos *α*] a unit vector defining the orientation of the fibres (*α* = 0, 90, ±*α*_0_ corresponds to axial, circumferential, and diagonal directions).

For each group of the particles (elastin, intra-lamellar, and adventitial), the total strain energy function has different mixing fractions of elastin and collagen fibres. We multiply the strain energy of each constituent (i.e. elastin and collagen families “*k*” denoted by superscripts *e* and *c*_*k*_) by an appropriate mass fraction, namely 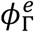 and 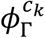, where Γ = *M*_*el*_, *M*_*int*_, *A* is used to distinguish elastin, intra-lamellar, and adventitial particles respectively. We use the total deformation gradient (***F***_*j*_) and the corresponding right Cauchy-Green (***C***_*j*_ = (***F***_*j*_)^T^***F***_*j*_) tensor to describe deformations from the homeostatic reference to any non-homeostatic configuration. To reproduce the stresses of the wall at the homeostatic reference configuration, we define deposition pre-stretches specific to each constituent in such a way that the resulting wall stress is in equilibrium with the homeostatic pressure and axial stretch. For a particle *j*, the deposition pre-stretch tensor for elastin is denoted by 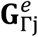 and deposition pre-stretches for the smooth muscle cells and collagen fibre family *k* are denoted 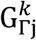, where Γ = *M*_*el*_, *M*_*int*_, *A* denotes the group of the particles (elastin, intra-lamellar, and adventitial particles). By following the numerical algorithm presented in (Ahmadzadeh et al., 2018), the deposition pre-stretches are introduced in the deformation gradient of elastin through 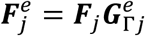, and the right Cauchy-Green tensor of collagen by 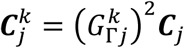.

The parameters embedded within the model were estimated using our previous analytical model of a normal aorta and fitting the predicted pressure-outer diameter and axial force-length results to the experimentally measured data for the DTA of the progeria mouse at 100 days of age. Because the analytical approach is based on a bilayered thin-walled vessel, the obtained parameters are adjusted to account for the lamellar structure of the media in this particle-based model. These parameters are presented in Table S8.

Finally, to introduce a swelling pressure caused by accumulated proteoglycans in the intra-lamellar regions, we randomly prescribed intra-lamellar particles (i.e., selected particles stochastically for 10%, 20%, or 30% GAGs within the media, noting that the different stochastic distributions had no effect on structural stiffness for a given mass fraction of GAGs) and modified their Cauchy stress function to

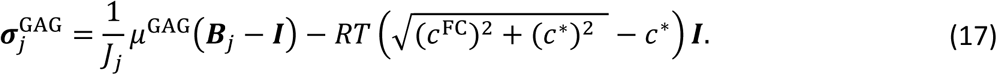

The first term represents the background stiffness of the proteoglycan pools (*μ*^GAG^ = 0.1 *kPa* as used previously) with the left Cauchy-Green tensor defined as 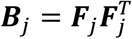. The second term accounts for the Gibbs-Donnan swelling pressure caused by the proteoglycans due to the absorption of water into the interstitial space occupied by the GAGs. *R* and *T* are the universal gas constant and the absolute body temperature (310 °K), *c*^*^ is the ionic concentration of the surrounding medium (assumed to be *c*^*^ = 300 mEq/l, consistent with prior work), and *c*^FC^ is the fixed charge density associated with the concentration of GAGs (with *c*^*^ = 200 mEq/l, equivalent to ∼155 kPa Gibbs-Donnan swelling pressure).

With regard to the specific simulations in Fig. 5, to account for the change in the homeostatic axial stretch from *λ*_*z*_ = 1.4 at 100 days to *λ*_*z*_ = 1.17 at 140 days, the axial stretch was reduced gradually from 1.40 to 1.17 to maintain quasi-static equilibrium. To account for different collagen pre-stretches, the adventitial value was increased by 5%, 10%, and 15% (e.g., from 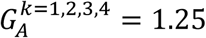 in Table S8 to 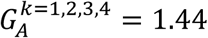). Similar steps were repeated on the baseline model when increasing the stiffness parameters of the adventitial collagen, for example, from 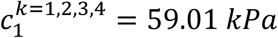 to 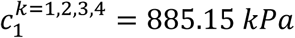 and from 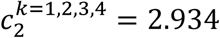 to 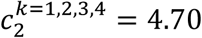 (cf. Table S8). We also repeated similar steps for an aorta but with normal collagen that simply doubled the adventitia thickness.

### Statistics

An analysis of variance (ANOVA) was used to compare results across region, genotype, or sex, with Bonferroni post-hoc testing and *p* < 0.05 considered significant.

## ACKNOWLEDGMENTS

We acknowledge expert advice on the *in vivo* data collection and analysis from Dr. Jacopo Ferruzzi, then at Yale University. This work was supported, in part, by grants from the US National Institutes of Health: R01 HL105297 (JDH) and P01 HL134605 (Dan Rifkin) and R01 AG047632 and R33 ES025636 (GSS).

## AUTHOR CONTRIBUTIONS

Designed Research: S-IM, DTB, JDH; Performed Research: S-IM, YK, AWC, HA, NM, KZ, DK, DW, ML, ZWZ; Analyzed Data: S-IM, YK, HA, DK, DTB, JDH; Contributed New Reagents or Tools: GSS; Wrote the Paper: S-IM, JDH.

## SUPPLEMENTAL TABLES

**Table S1.**
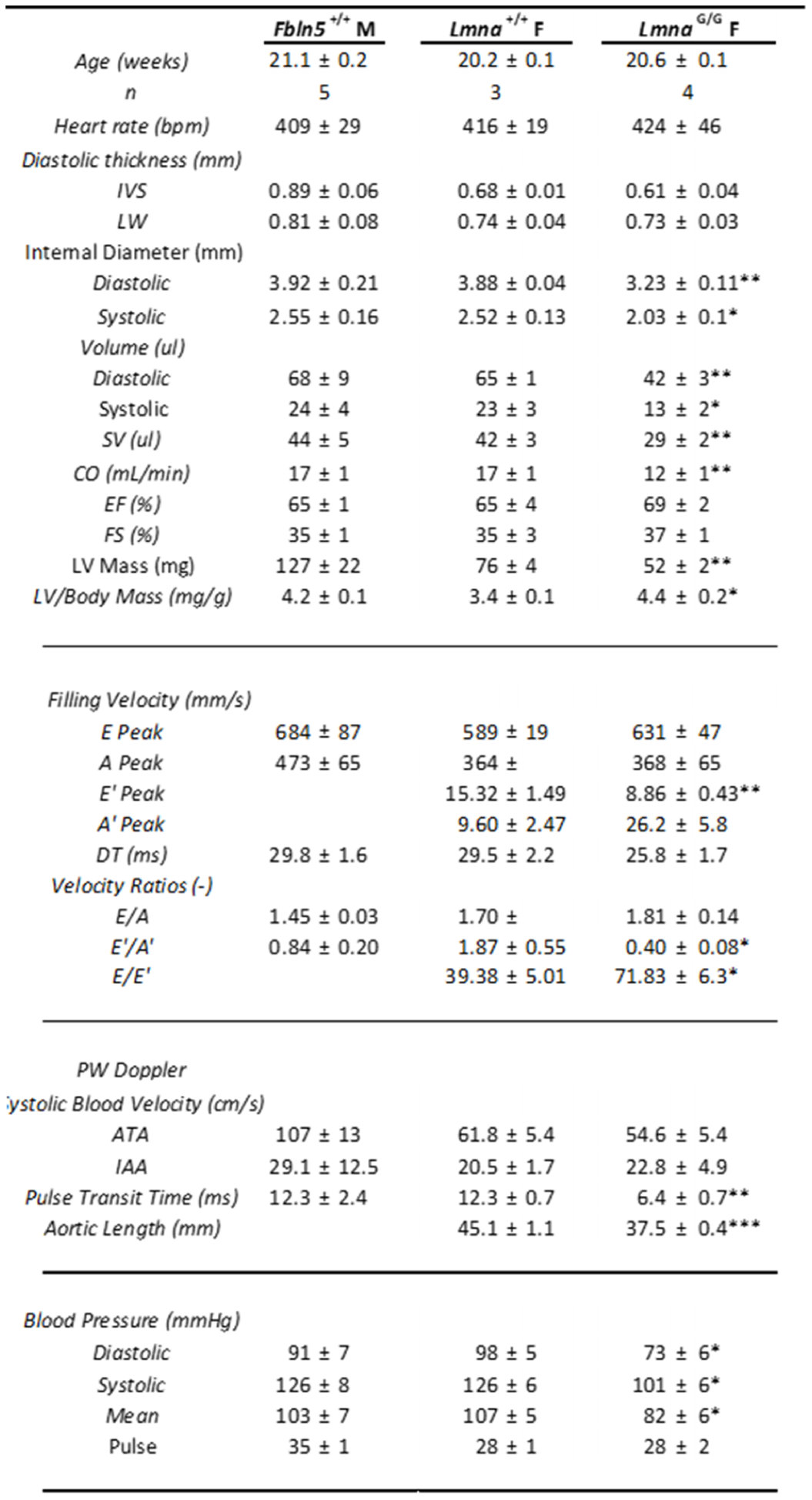
Echocardiographic data plus ultrasonic data for central arteries for age-matched (140-day old) wild-type controls (*Fbln5*^*+/+*^; Ferruzzi et al., 2018) as well as littermate (*Lmna*^*+/+*^) and progeria (*Lmna*^*G/G*^ = *Lmna*^G609G/G609G^) mice, plus conscious tail-cuff measured blood pressures. Note that tail-cuff pressures enable mouse-to-mouse comparisons of most biomechanical metrics but they need not mimic central arterial pressures, which can be elevated due to the earlier wave reflections due to increased PWV (Cuomo et al., 2019). ***: P<0.001, **: P<0.01, *: P<0.05.

**Table S2.** Geometrical and mechanical metrics (mean ± SEM) for the contractile behavior of the aorta, by region (DTA – descending thoracic aorta, and IAA – infrarenal thoracic aorta), for mixed-sex (F plus M) littermate control (*Lmna*^+/+^) and progeria (*Lmna*^G609G/G609G^) mice at 100 days of age. ***: P<0.001, **: P<0.01, *: P<0.05.

| 100 days |  | DTA |  | IAA |  |
| --- | --- | --- | --- | --- | --- |
|  |  | *Lmna*<sup>+/+</sup><br>n = 5 | *Lmna*<sup>G/G</sup><br>n = 5 | *Lmna*<sup>+/+</sup><br>n = 5 | *Lmna*<sup>G/G</sup><br>n = 5 |
| 100 mM KCl | **Loaded configuration** |  |  |  |  |
|  | Relaxed Outer Diameter (μm) | 1103 ± 24 | 955 ± 21*** | 768 ± 20 | 648 ± 11*** |
|  | Contracted Outer Diameter (μm) | 915 ± 35 | 838 ± 23 | 675 ± 21 | 589 ± 25* |
|  | Change in Outer Diameter (%) | -17 ± 2.0 | -12 ± 1.8 | -12 ± 1.9 | -9 ± 3.1 |
|  | **Stress calculations** |  |  |  |  |
|  | Relaxed Circumferential Stress (kPa) | 145 ± 4.4 | 95 ± 4.3*** | 123 ± 14.2 | 80 ± 7.8* |
|  | Contracted Circumferential Stress (kPa) | 95 ± 7.7 | 69 ± 1.7* | 91 ± 13.4 | 64 ± 10 |
|  | Change in Circumferential Stress (%) | -35 ± 3.9 | -27 ± 3.6 | -26 ± 4.0 | -21 ± 6.8 |
|  | Active Circumferential Force (mN) | 21 ± 2.9 | 10 ± 1.4** | 11 ± 2.5 | 7 ± 3.2 |
| 1 μM PE | **Loaded configuration** |  |  |  |  |
|  | Relaxed Outer Diameter (μm) | 1120 ± 31 | 960 ± 26** | 762 ± 34 | 663 ± 11* |
|  | Contracted Outer Diameter (μm) | 878 ± 24 | 869 ± 19 | 585 ± 38 | 585 ± 26 |
|  | Change in Outer Diameter (%) | -22 ± 0.9 | -9 ± 2.1*** | -23 ± 4.7 | -12 ± 3 |
|  | **Stress calculations** |  |  |  |  |
|  | Relaxed Circumferential Stress (kPa) | 150 ± 4.2 | 96 ± 3.3*** | 122 ± 17.3 | 85 ± 6.8 |
|  | Contracted Circumferential Stress (kPa) | 85 ± 4.6 | 75 ± 2.9 | 66 ± 15.1 | 63 ± 9.9 |
|  | Change in Circumferential Stress (%) | -43 ± 1.8 | -22 ± 4.3** | -47 ± 9.4 | -27 ± 6.7 |
|  | Active Circumferential Force (mN) | 28 ± 2.4 | 8 ± 1.8*** | 24 ± 6.1 | 8 ± 2.8 |
|  | **Metrics normalized to KCl** |  |  |  |  |
|  | Change in Diameter (%) | 137 ± 20 | 90 ± 28 | 209 ± 52 | 146 ± 15 |

**Table S3.**
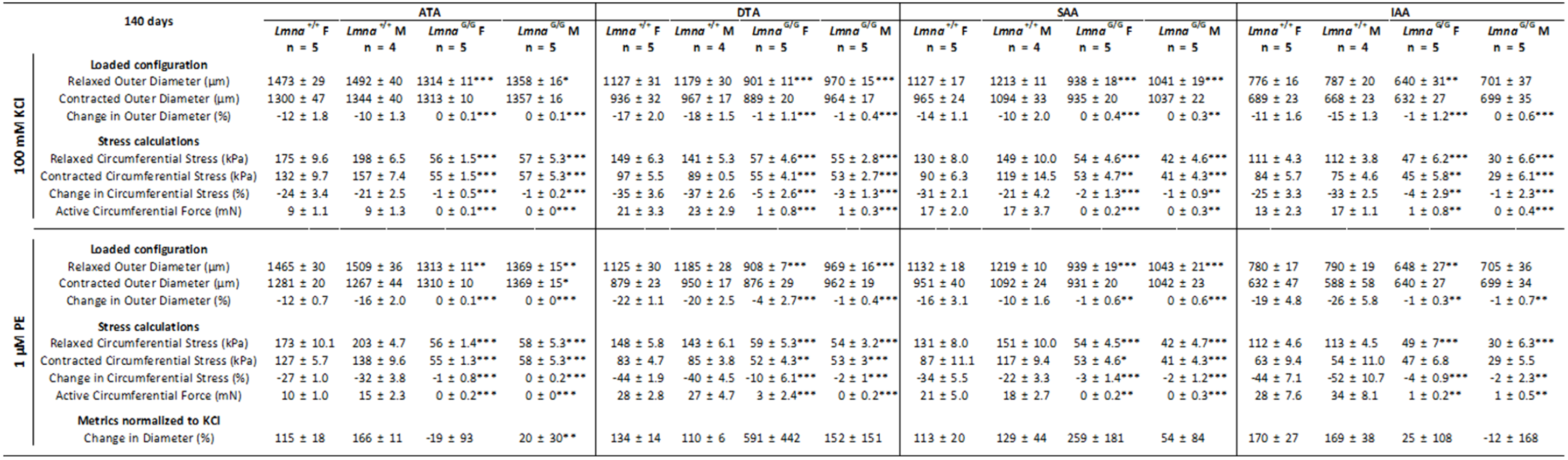
Geometrical and mechanical metrics (mean ± SEM) for the contractile behavior of the aorta, by region (ATA – ascending thoracic aorta, DTA – descending thoracic aorta, SAA – suprarenal thoracic aorta, and IAA – infrarenal thoracic aorta), for both male (M) and female (F) littermate control (*Lmna*^+/+^) and progeria (*Lmna*^G609G/G609G^) mice at 140 days of age. Note the dramatic loss of contractile function from 100d to 140d, consistent with the overall worsening of the aortic phenotype and cardiac diastolic function. ***: P<0.001, **: P<0.01, *: P<0.05.

**Table S4.** Best-fit values of the material parameters in the nonlinear constitutive relation (see Methods) for the passive aortic behavior at 100 days, by region (DTA – descending thoracic aorta, and IAA – infrarenal thoracic aorta), for mixed-sex (F plus M) mice for both genotypes (*Lmna*^*+/+*^ and *Lmna*^*G/G*^ = *Lmna*^*G609G/G609G*^). Although not shown, the quantitative fit to the biaxial biomechanical data was similar to that achieved previously in multiple mouse models (Ferruzzi et al., 2015; Bersi et al., 2016; Bellini et al., 2017), noting that the associated “four-fiber family” stored energy function has also been independently validated by multiple groups. ***: P<0.001, **: P<0.01, *: P<0.05.

| 100 days |  | Elastic fibers | Axial Collagen |  | Circumf. Coll +SMC |  | Symmetric diagonal collagen |  |  | Error |
| --- | --- | --- | --- | --- | --- | --- | --- | --- | --- | --- |
| | | $c$ (kPa) | $c_1^1$ (kPa) | $c_2^1$ | $c_1^2$ (kPa) | $c_2^2$ | $c_1^{3,4}$ (kPa) | $c_2^{3,4}$ | $\alpha_o$ (deg) | RMSE |
| DTA | *Lmna*<sup>+/+</sup> | 30.235 | 10.081 | 0.18 | 11.797 | 0.064 | 1.308 | 0.977 | 32.8 | 0.0692 |
|  | *Lmna*<sup>G/G</sup> | 19.793 | 12.136 | 0.68 | 15.05 | 0.086 | 2.729 | 1.668 | 27.3* | 0.0763 |
| IAA | *Lmna*<sup>+/+</sup> | 16.853 | 4.551 | 0.23 | 8.687 | 0.256 | 0.738 | 0.676 | 43.2 | 0.0959 |
|  | *Lmna*<sup>G/G</sup> | 16.297 | 3.759 | 0.46 | 8.804 | 0.464 | 1.647 | 1.371 | 41.4 | 0.1070 |

**Table S5.**
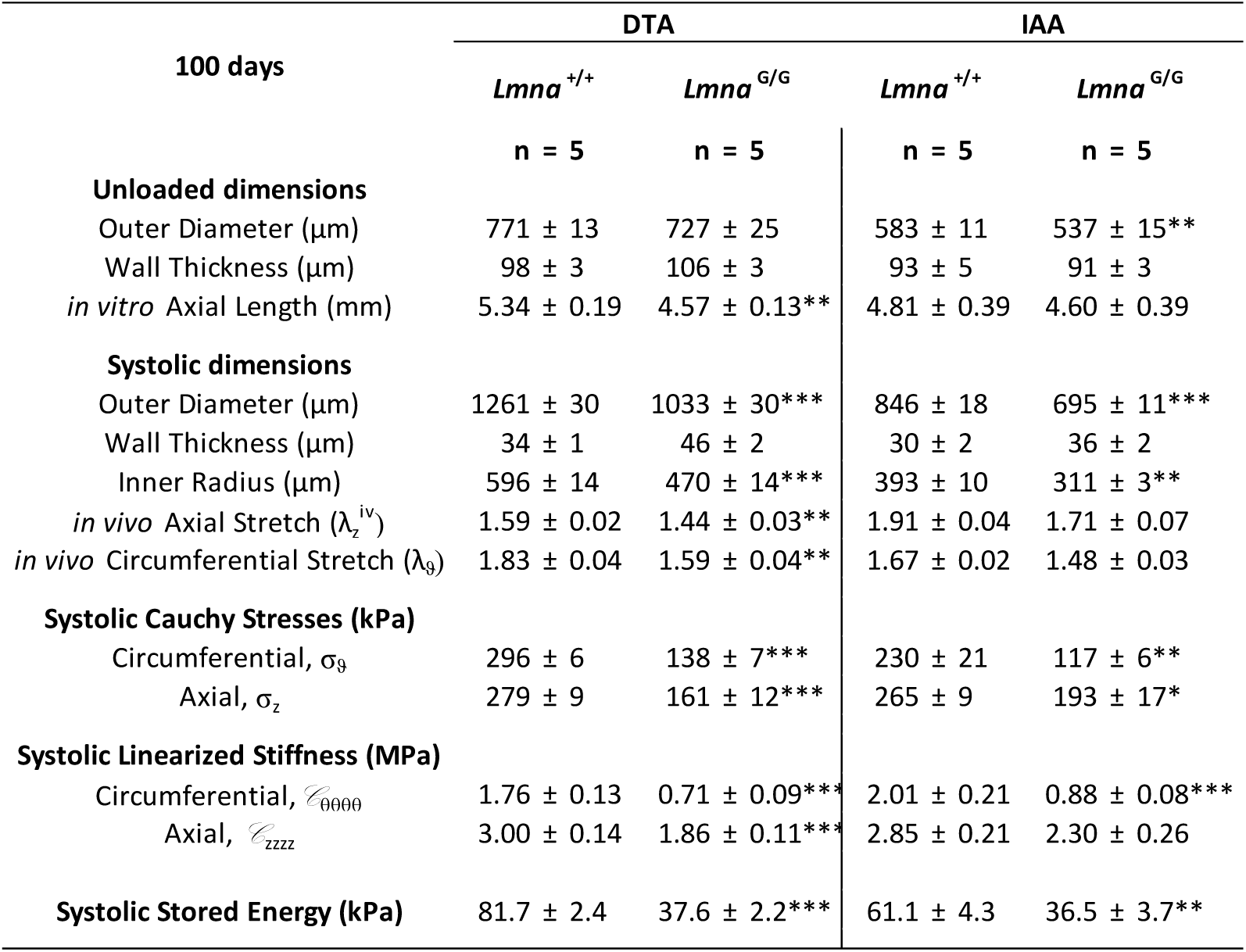
Geometrical and mechanical metrics (mean ± SEM) for the passive behavior of the aorta, by region (DTA – descending thoracic aorta, and IAA – infrarenal thoracic aorta), for mixed-sex (F plus M) littermate control (*Lmna*^+/+^) and progeria (*Lmna*^G609G/G609G^) mice at 100 days of age. Computed values are based on group-specific pressures and mouse-specific in vivo values of axial stretch, which enable consistent comparisons. ***: P<0.001, **: P<0.01, *: P<0.05.

**Table S6.** Best-fit values of the material parameters in the nonlinear constitutive relation for the passive behavior at 140 days, by aortic region (ATA – ascending thoracic aorta, DTA – descending thoracic aorta, SAA – suprarenal thoracic aorta, and IAA – infrarenal thoracic aorta), for both sexes (F and M) and genotypes (*Lmna*^*+/+*^ and *Lmna*^*G/G*^ = *Lmna*^*G609G/G609G*^). See also Table S4. ***: P<0.001, **: P<0.01, *: P<0.05.

| 140 days |  | Elastic fibers | Axial Collagen |  | Circumf. Coll +SMC |  | Symmetric diagonal collagen |  |  | Error |
| --- | --- | --- | --- | --- | --- | --- | --- | --- | --- | --- |
| | | $c$ (kPa) | $c_1^{-1}$ (kPa) | $c_2^{-1}$ | $c_1^{-2}$ (kPa) | $c_2^{-2}$ | $c_1^{-3,4}$ (kPa) | $c_2^{-3,4}$ | $\alpha_o$ (deg) | RMSE |
| ATA | *Lmna*<sup>+/+</sup> F | 20.586 | 7.917 | 0.03 | 8.081 | 0.709 | 8.966 | 0.242 | 49.3 | 0.0799 |
|  | *Lmna*<sup>+/+</sup> M | 24.676 | 11.284 | 0.03 | 9.166 | 0.759 | 9.623 | 0.252 | 47.5 | 0.0818 |
|  | *Lmna*<sup>G/G</sup> F | 12.466 | 4.009 | 12.11 | 18.642 | 1.775 | 26.629* | 4.244*** | 39.8** | 0.0742 |
|  | *Lmna*<sup>G/G</sup> M | 18.065 | 10.943 | 10.79** | 11.084 | 2.319 | 18.109 | 5.533* | 40.9 | 0.0929 |
| DTA | *Lmna*<sup>+/+</sup> F | 25.647 | 7.939 | 0.25 | 10.164 | 0.075 | 0.949 | 0.938 | 34.1 | 0.0827 |
|  | *Lmna*<sup>+/+</sup> M | 21.551 | 7.072 | 0.27 | 11.750 | 0.041 | 1.347 | 0.861 | 35.3 | 0.0861 |
|  | *Lmna*<sup>G/G</sup> F | 19.947 | 11.248 | 8.67* | 14.687 | 0.425* | 5.943* | 6.25** | 27.7* | 0.0642 |
|  | *Lmna*<sup>G/G</sup> M | 26.974 | 5.373 | 14.91* | 21.726 | 3.602 | 9.094 | 17.604* | 28.4 | 0.0891 |
| SAA | *Lmna*<sup>+/+</sup> F | 23.123 | 5.406 | 0.25 | 7.723 | 0.190 | 0.890 | 1.028 | 40.9 | 0.0791 |
|  | *Lmna*<sup>+/+</sup> M | 24.585 | 5.045 | 0.23 | 7.617 | 0.165 | 0.669 | 0.982 | 38.1 | 0.0818 |
|  | *Lmna*<sup>G/G</sup> F | 22.113*** | 8.143*** | 5.46*** | 7.163*** | 3.609*** | 3.411*** | 9.013*** | 37.2*** | 0.0533 |
|  | *Lmna*<sup>G/G</sup> M | 17.929 | 1.458 | 8.52*** | 5.801 | 17.059 | 4.143* | 15.454*** | 46.2 | 0.0973 |
| IAA | *Lmna*<sup>+/+</sup> F | 17.693 | 2.431 | 0.28 | 7.856 | 0.409 | 0.232 | 1.001 | 41.5 | 0.0979 |
|  | *Lmna*<sup>+/+</sup> M | 17.072 | 2.446 | 0.26 | 5.962 | 0.426 | 0.359 | 0.914 | 41.0 | 0.0938 |
|  | *Lmna*<sup>G/G</sup> F | 14.717*** | 3.111*** | 2.23*** | 8.301*** | 1.142*** | 0.545*** | 4.116*** | 38*** | 0.0853 |
|  | *Lmna*<sup>G/G</sup> M | 9.469 | 1.886 | 8.25 | 12.27 | 13.413* | 12.296 | 12.876* | 49.1 | 0.1811 |

**Table S7.**
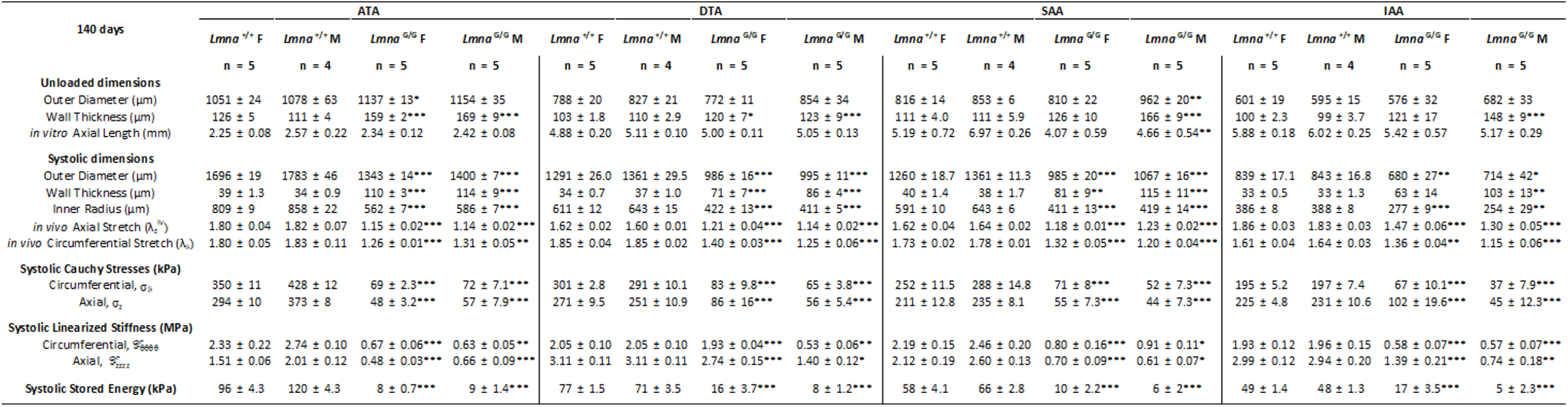
Geometrical and mechanical metrics (mean ± SEM) for the passive behavior of the aorta, by region (ATA – ascending thoracic aorta, DTA – descending thoracic aorta, SAA – suprarenal thoracic aorta, and IAA – infrarenal thoracic aorta), for both male (M) and female (F) littermate control (*Lmna*^+/+^) and progeria (*Lmna*^G609G/G609G^) mice at 140 days of age. See also Table S5. ***: P<0.001, **: P<0.01, *: P<0.05.

**Table S8.**
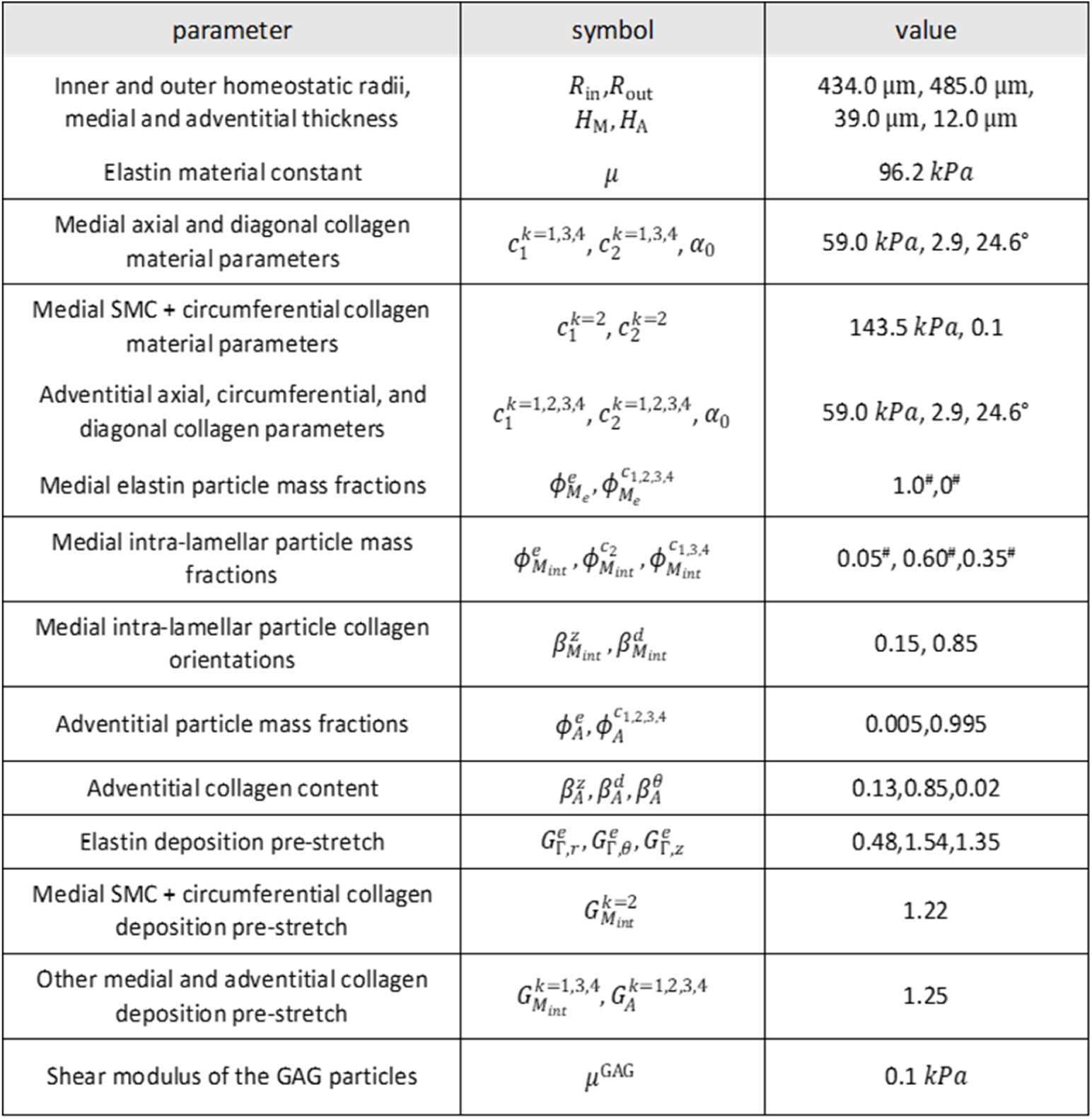
Best-fit parameters for the multi-layered, particle-based computational model. See Methods for the specific equations and methods of solution. # denotes that the fitted parameter was adjusted to account for the lamellar structure of the media, which has not been considered in prior analytical solutions. In particular, this model is the first to allow one to delineate separate mechanical contributions by the elastic lamellar structures, the passive and active smooth muscle cells, the medial and adventitial collagen fibres, and the presence of stochastically distributed glycosaminoglycans (GAGs). Consistent with an earlier analysis of medial GAGs in different conditions (Roccabianca et al., 2014), we found that the Gibbs-Donnan swelling pressures associated with the highly negatively charged GAGs were largely responsible for widening of the intra-lamellar spaces, which in turn would be expected to overly stress physical connections between the smooth muscle cells and the elastic laminae, with expected compromised mechano-sensing at the minimum and anoikis-induced cell death in the extreme (cf. Meredith et al., 1993). Finally, the simulations showed that it is not the amount of collagen, but rather its altered mechanical properties that contributes to the observed aortic behavior.

## SUPPLEMENTAL FIGURES

**Fig. S1.**
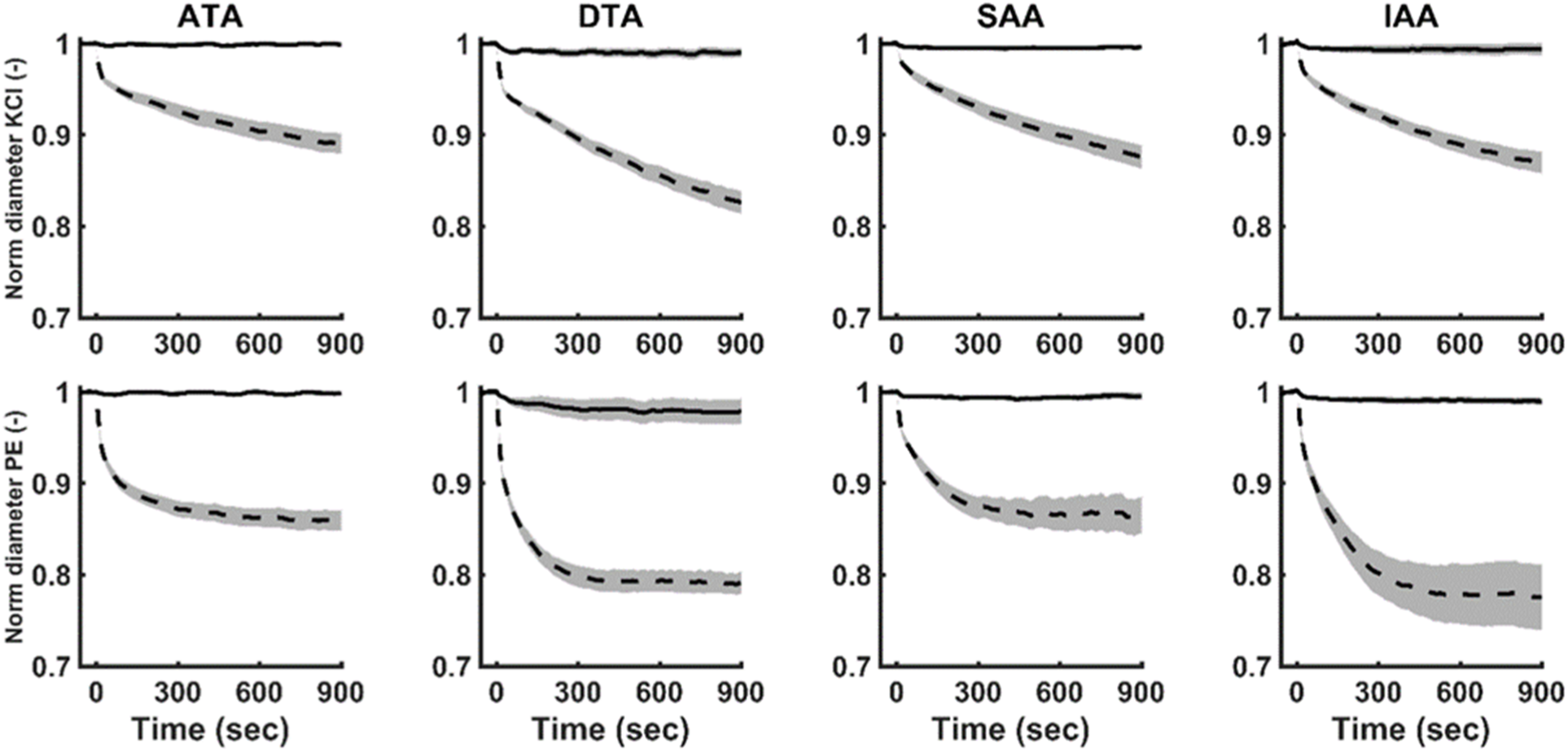
Similar to Figure 2 in the text, but showing that all four aortic regions (ATA,DTA,SAA,IAA) are adversely affected at 140 days of age. Specifically, we show normalized reduction in outer diameter during *ex vivo* biaxial (isobaric-isometric) active contraction in response to (**A-D**) 100 mM KCl or (**E-H**) 1 μM phenylephrine, both at 90 mmHg and group-specific values of *in vivo* axial stretch from mixed-sex *Wt* (dashed lines) and *G609G* (solid curve) mice.

**Fig. S2.**
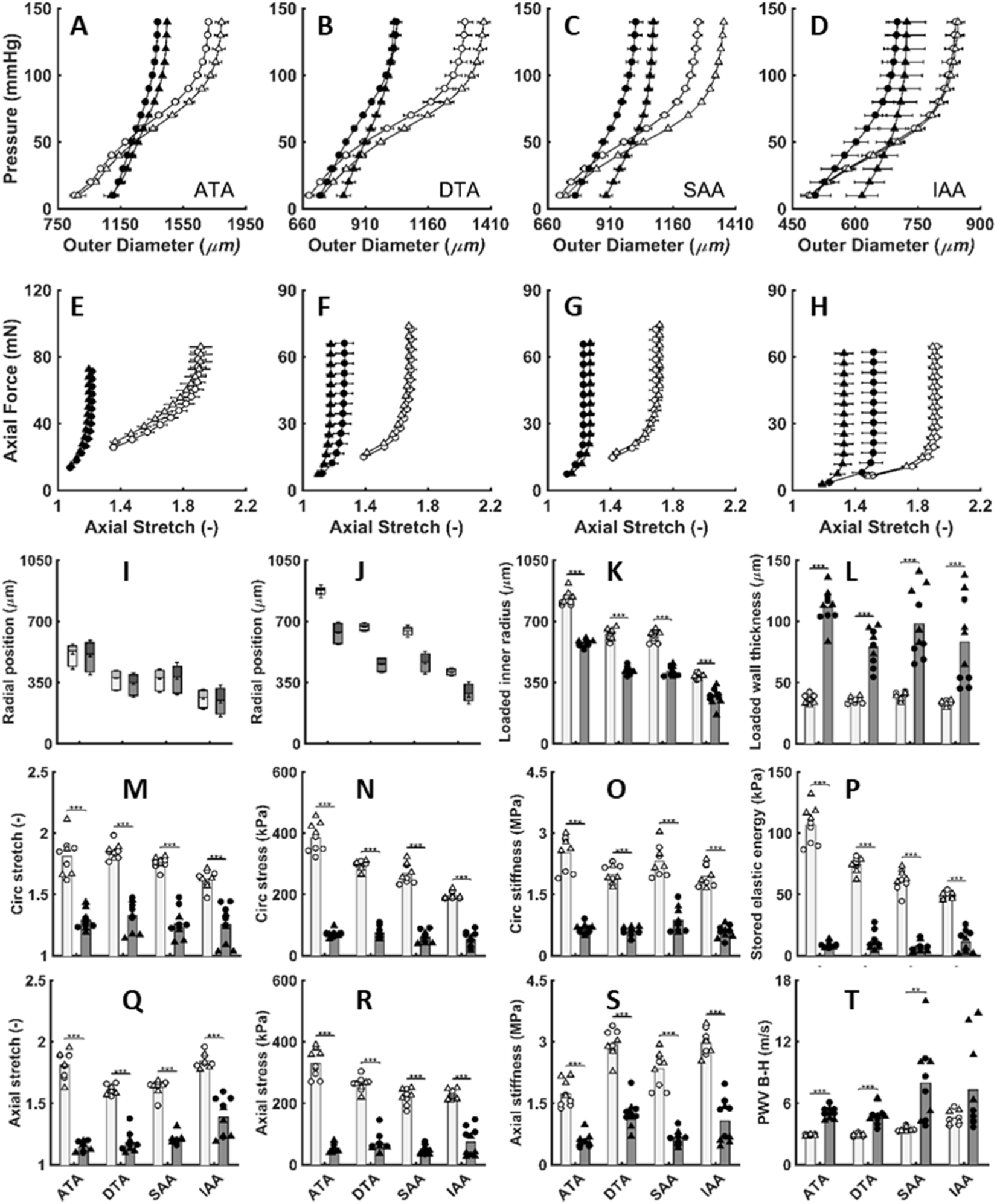
Similar to Figure 3 in the text, but showing that all four aortic regions (ATA,DTA,SAA,IAA) are adversely affected at 140 days of age. Specifically, **A-H:** Pressure–Outer diameter and Axial force – Stretch relationships of four regions from 140-day-old female (o) and male (Δ) *Wt* (open) and *G609G (closed)* mice. **I-T:** Computed geometrical and mechanical metrics from the passive tests on the 140-day-old female (o) and male (Δ) control (open) and progeria (filled) aortas under in vivo (systolic) conditions. (**I-L**) Unloaded and loaded radial position of medial and adventitial layer, inner radius, and wall thickness. (**M,Q**) Biaxial stretch, (**N,R**) stress, and (**O,S**) intrinsic material stiffness. (**P)** Stored elastic energy *W* and (**L**) Bramwell-Hill pulse wave velocity PWV. Note that the segmental PWVs are all elevated, but especially so in the more distal aorta where there was also greater variability. Natural aging similarly tends to affect the distal (abdominal) aorta more than the proximal (thoracic) aorta (Ferruzzi et al.,, 2018b). ***: P<0.001, **: P<0.01, *: P<0.05.

**Fig. S3.**
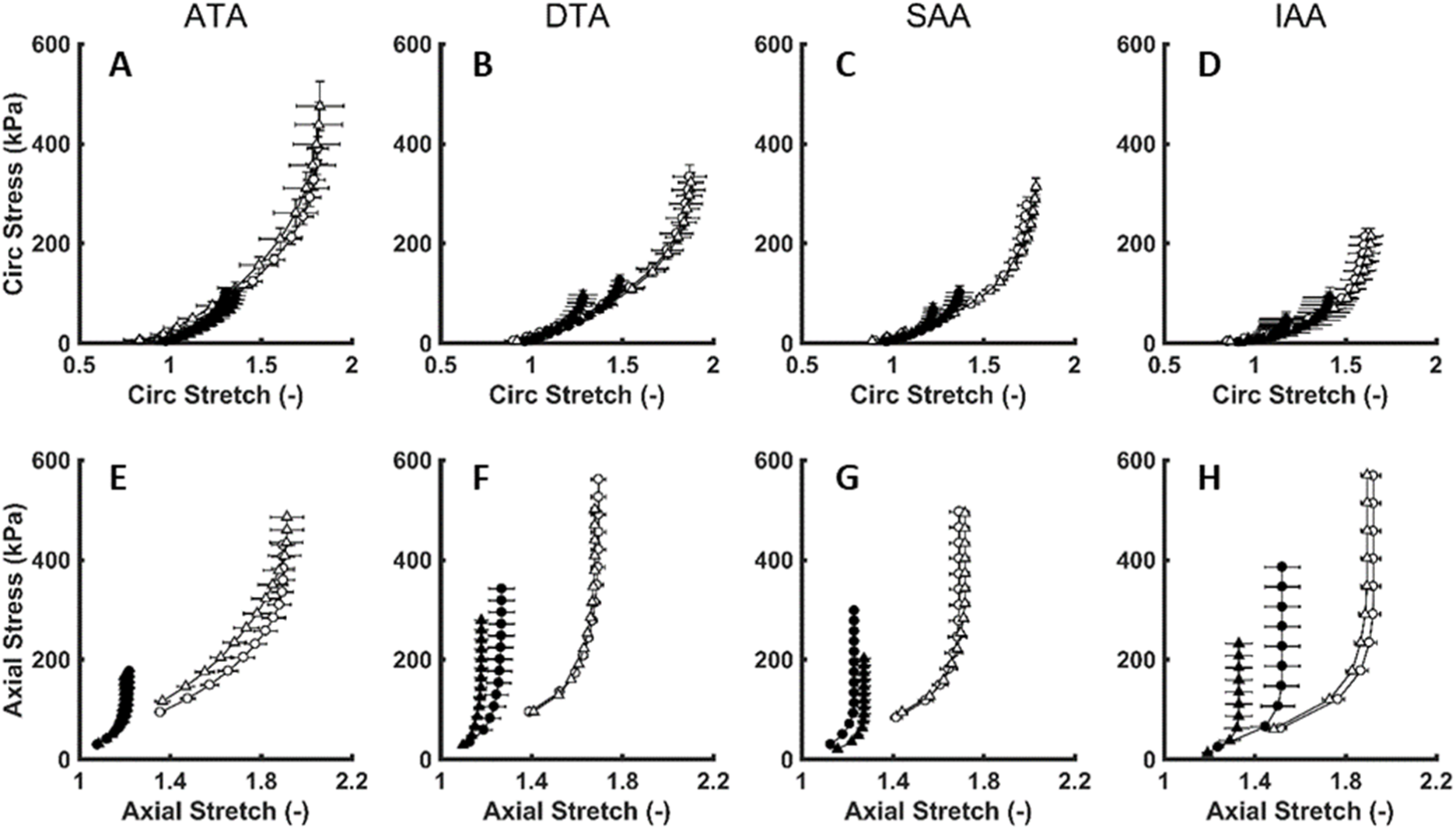
(**A-D**) Circumferential and (**E-H**) axial stress-stretch relationships of four aortic regions (ATA,DTA,SAA,IAA) from female (o) and male (Δ) 140-day-old Wt (open) and G609G (solid) mice. Although differences are readily seen (cf. del Campo et al., 2018), such standard stress-strain plots typically provide less information than detailed calculations of material stiffness and energy, which are provided in Figures 3 and S2 and Tables S5 and S7.

**Fig. S4.**
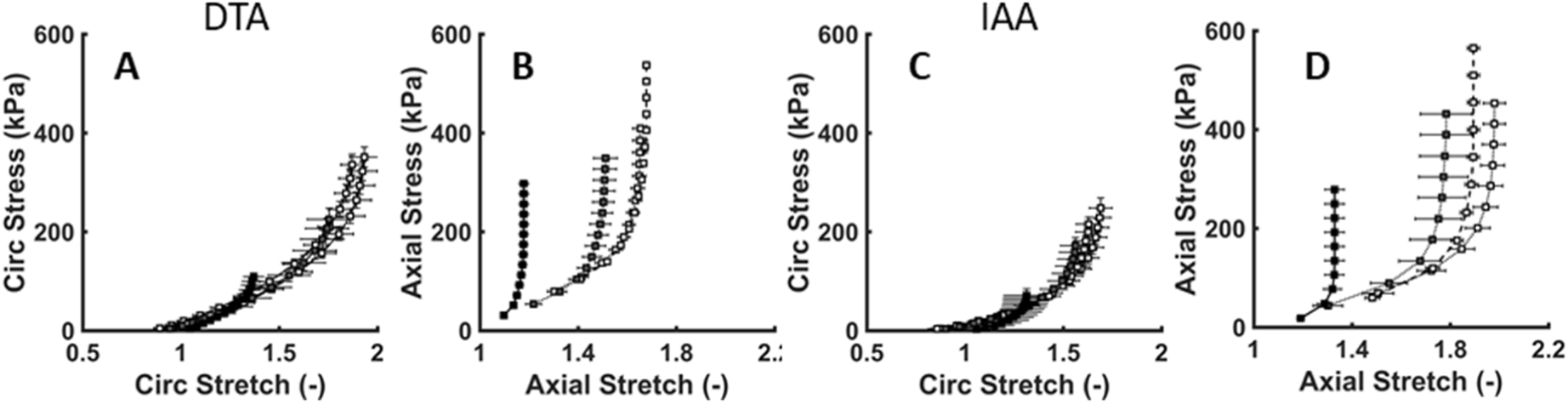
(**A,C**) Circumferential and (**B,D**) axial stress-stretch relationships for DTA and IAA from mixed sex (□) 100- (grey) and 140-day-old (black) Wt (open) and G609G (filled) mice.

**Figure S5.**
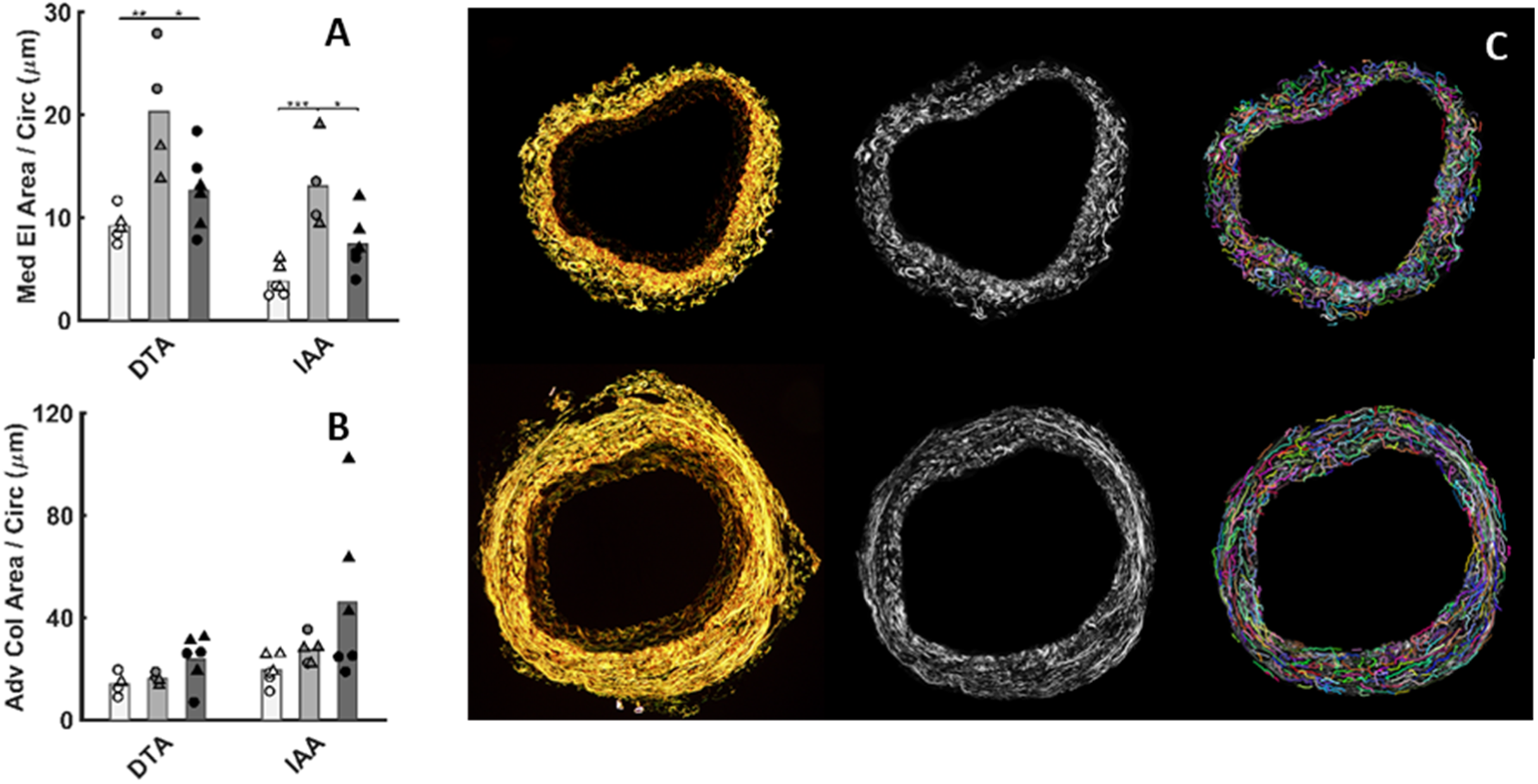
Quantification of (**A**) medial elastin (VVG) and (**B**) adventitial collagen (MOV), each normalized by individual specimen inner vessel circumference at systolic loading conditions to account for different aortic size by region, sex, body size, age, and genotype. An increase in elastin concentration (Fig. S5A) was observed in the media between *Wt* and *G609G* aorta, likely due to an allometric effect, but no significant difference was observed in adventitial collagen. Estimation of (**C**) adventitial collagen fiber undulation using fibre-tracking algorithm from gray scaled histological PSR images (Bredfeldt et al., 2014), noting that the computational model provided the best information on the associated collagen properties (Figure 5).

**Figure S6.**
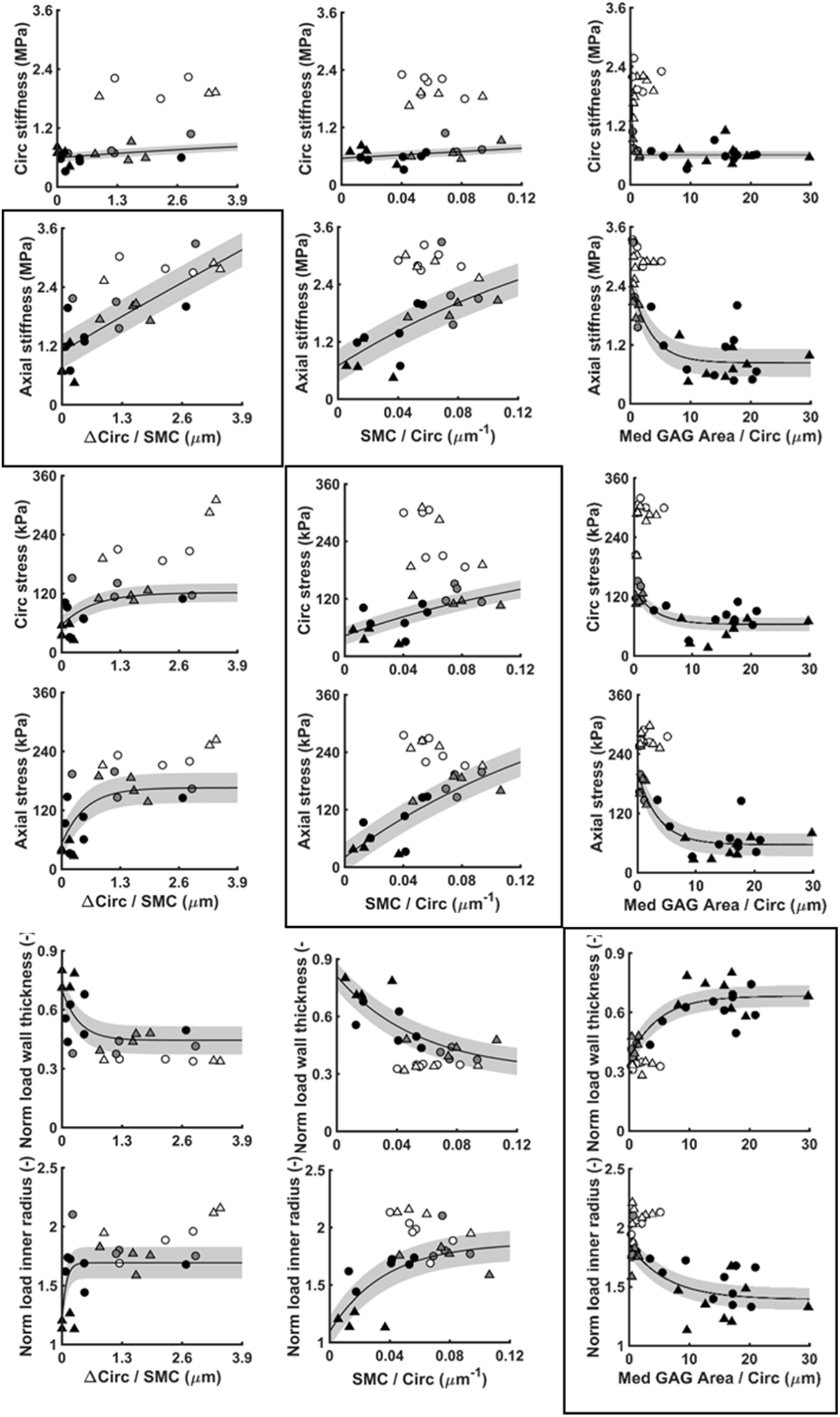
Computed vascular geometrical and mechanical metrics from passive tests on the 100- and 140-day-old female (o) and male (Δ) *Wt* (open) and *G609G* (filled) aortas under *in vivo* (systolic) conditions. Circumferential and axial (**A-F)** material stiffness and (**G-L**) stress, and (**M-S)** loaded wall thickness and inner radius, shown as a function of SMC contractile response (change in circumference per SMC) as well as SMC density normalized by loaded per circumference and medial GAG per circumference, all at systole. While circumferential stiffness seems to be independent to SMC contractile response, SMC density, and medial GAG, correlations emerged among particular mechanical metrics (five boxed figures).

**Figure S7.**
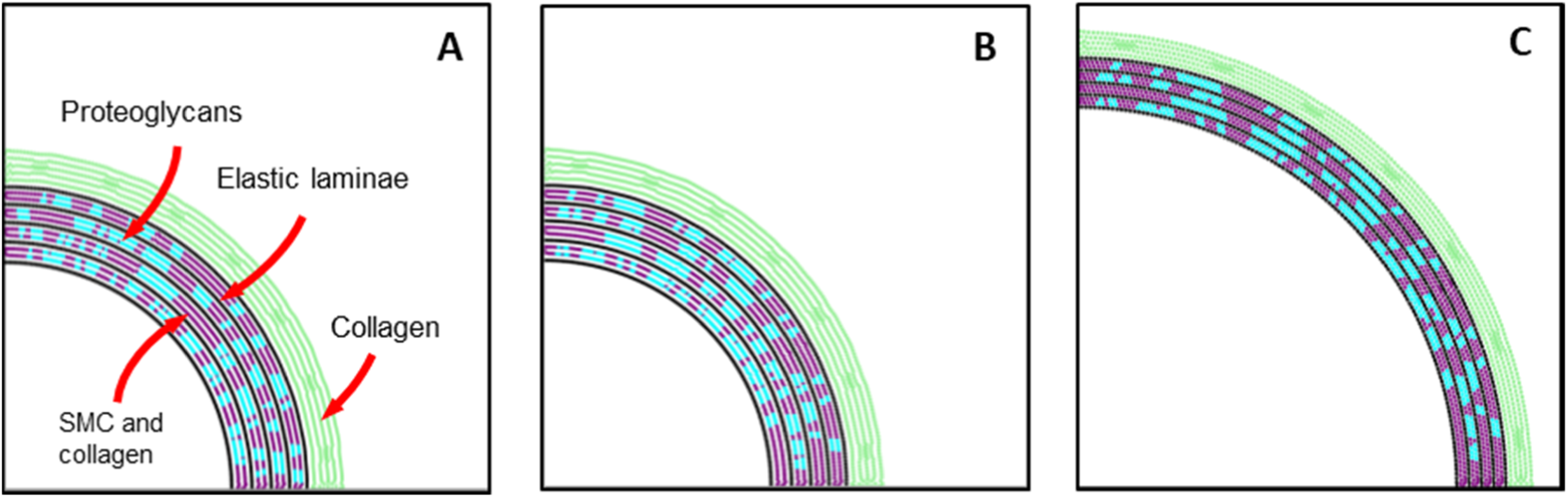
To verify the independence of the particle-based computational results from the distribution of the pools of proteoglycans, four different stochastic distributions, two of which are shown in **(A)** and **(B)**, were tested and the associated pressure-outer diameter relations are shown in Fig. 5F. While the wall in **(A)** and **(B)** are at the zero-pressure state for comparison, the pressurized state of the wall is also shown in **(C)**.

**Figure S8.**
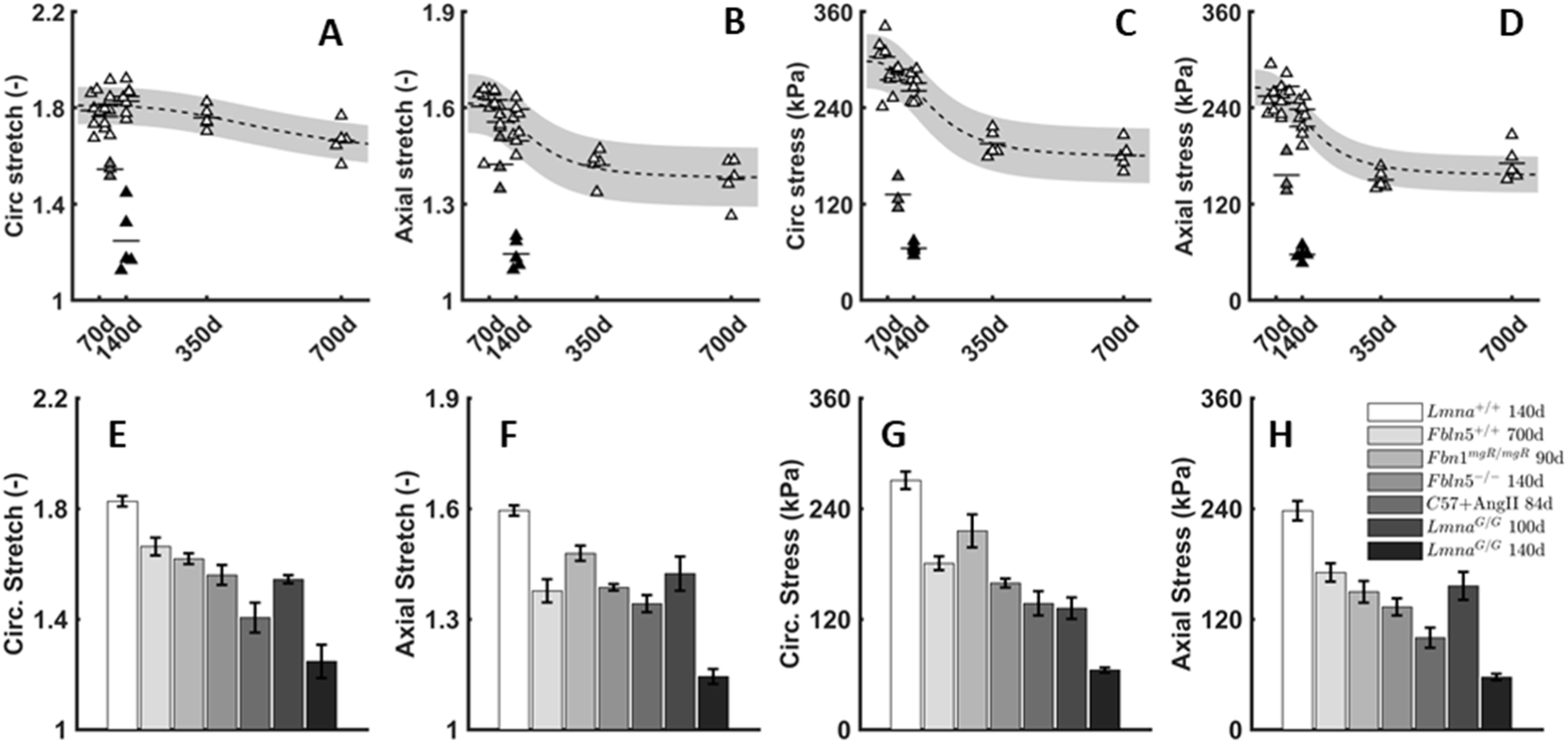
Circumferential (**A,E**) and axial (**B,F**) stretch and (**C,G,D,H**) wall stress for the descending thoracic aorta, as presented in Fig. 6 for male mice, as a function of (**A-D**) natural aging between 56d-700d of age and (**E-H**) aging-related disease conditions: naturally aged to 700d, fibrillin-1 deficient (*Fbln1*^*mgR/mgR*^), fibulin-5 null (*Fbln5*^*-/-*^), or hypertensive (C57+AngII) are compared with 100d and 140d old *G609G* mice, each calculated at group-specific systolic pressures and *in vivo* axial stretch. Again it is appreciated that the 100d-old progeria aorta is similar to that of the 100-wk-old naturally aged aorta, thus revealing the extent of the pre-mature aging. Yet, the 140d-old progeria aorta again shows the most extreme phenotype of all conditions considered, emphasizing again the rapid worsening in the late-stage progeria aorta that associates with the appearance of the glycosaminoglycans and complete loss of vasoconstrictive capacity.

## REFERENCES

Ahmadzadeh, H., Rausch, M. K., & Humphrey, J. D. (2018). Particle-based computational modelling of arterial disease. Journal of the Royal Society Interface, 15(149), 20180616. https://doi.org/10.1098/rsif.2018.0616

Bellini, C., Bersi, M. R., Caulk, A. W., Ferruzzi, J., Milewicz, D. M., Ramirez, F., … & Humphrey, J. D. (2017). Comparison of 10 murine models reveals a distinct biomechanical phenotype in thoracic aortic aneurysms. Journal of The Royal Society Interface, 14(130), 20161036. https://doi.org/10.1098/rsif.2016.1036

Bellini, C., Ferruzzi, J., Roccabianca, S., Di Martino, E. S., & Humphrey, J. D. (2014). A microstructurally motivated model of arterial wall mechanics with mechanobiological implications. Annals of biomedical engineering, 42(3), 488–502. https://doi.org/10.1007/s10439-013-0928-x

Bersi, M. R., Bellini, C., Wu, J., Montaniel, K. R., Harrison, D. G., & Humphrey, J. D. (2016). Excessive adventitial remodeling leads to early aortic maladaptation in angiotensin-induced hypertension. Hypertension, 67(5), 890–896. https://doi.org/10.1161/HYPERTENSIONAHA.115.06262

Bredfeldt, J. S., Liu, Y., Pehlke, C. A., Conklin, M. W., Szulczewski, J. M., Inman, D. R., … & Eliceiri, K. W. (2014). Computational segmentation of collagen fibers from second-harmonic generation images of breast cancer. Journal of biomedical optics, 19(1), 016007. https://doi.org/10.1117/1.JBO.19.1.016007

Capell, B. C., Olive, M., Erdos, M. R., Cao, K., Faddah, D. A., Tavarez, U. L., … & Chen, X. (2008). A farnesyltransferase inhibitor prevents both the onset and late progression of cardiovascular disease in a progeria mouse model. Proceedings of the National Academy of Sciences, 105(41), 15902–15907. https://doi.org/10.1073/pnas.0807840105

Cikach, F. S., Koch, C. D., Mead, T. J., Galatioto, J., Willard, B. B., Emerton, K. B., … & Apte, S. S. (2018). Massive aggrecan and versican accumulation in thoracic aortic aneurysm and dissection. JCI insight, 3(5). https://doi.org/10.1172/jci.insight.97167

Cuomo, F., Ferruzzi, J., Agarwal, P., Li, C., Zhuang, Z. W., Humphrey, J. D., & Figueroa, C. A. (2019). Sex-dependent differences in central artery haemodynamics in normal and fibulin-5 deficient mice: implications for ageing. Proceedings of the Royal Society A, 475(2221), 20180076. https://doi.org/10.1098/rspa.2018.0076

Dajnowiec, D., & Langille, B. L. (2007). Arterial adaptations to chronic changes in haemodynamic function: coupling vasomotor tone to structural remodelling. Clinical Science, 113(1), 15–23. DOI: 10.1042/CS20060337

de Simone, G., Devereux, R. B., Daniels, S. R., Mureddu, G., Roman, M. J., Kimball, T. R., … & Contaldo, F. (1997). Stroke volume and cardiac output in normotensive children and adults: assessment of relations with body size and impact of overweight. Circulation, 95(7), 1837–1843. https://doi.org/10.1161/01.CIR.95.7.1837

del Campo, L., Sánchez-López, A., Salaices, M., von Kleeck, R. A., Expósito, E., González-Gómez, C., … & Assoian, R. K. (2019). Vascular smooth muscle cell-specific progerin expression in a mouse model of Hutchinson–Gilford progeria syndrome promotes arterial stiffness: Therapeutic effect of dietary nitrite. Aging cell, 18(3), e12936. https://doi.org/10.1111/acel.12936

Dewey, F. E., Rosenthal, D., Murphy Jr, D. J., Froelicher, V. F., & Ashley, E. A. (2008). Does size matter? Clinical applications of scaling cardiac size and function for body size. Circulation, 117(17), 2279–2287. https://doi.org/10.1161/CIRCULATIONAHA.107.736785

DuBose, A. J., Lichtenstein, S. T., Petrash, N. M., Erdos, M. R., Gordon, L. B., & Collins, F. S. (2018). Everolimus rescues multiple cellular defects in laminopathy-patient fibroblasts. Proceedings of the National Academy of Sciences, 115(16), 4206–4211. https://doi.org/10.1161/CIRCULATIONAHA.107.736785

Eberth, J. F., Taucer, A. I., Wilson, E., & Humphrey, J. D. (2009). Mechanics of carotid arteries in a mouse model of Marfan syndrome. Annals of biomedical engineering, 37(6), 1093–1104. https://doi.org/10.1007/s10439-009-9686-1

Eriksson, M., Brown, W. T., Gordon, L. B., Glynn, M. W., Singer, J., Scott, L., … & Dutra, A. (2003). Recurrent de novo point mutations in lamin A cause Hutchinson–Gilford progeria syndrome. Nature, 423(6937), 293. https://doi.org/10.1038/nature01629

Ferruzzi, J., Bersi, M. R., Uman, S., Yanagisawa, H., & Humphrey, J. D. (2015). Decreased elastic energy storage, not increased material stiffness, characterizes central artery dysfunction in fibulin-5 deficiency independent of sex. Journal of biomechanical engineering, 137(3), 031007. https://doi.org/10.1115/1.4029431

Ferruzzi, J., Di Achille, P., Tellides, G., & Humphrey, J. D. (2018). Combining in vivo and in vitro biomechanical data reveals key roles of perivascular tethering in central artery function. PloS one, 13(9), e0201379. https://doi.org/10.1371/journal.pone.0201379

Ferruzzi, J., Madziva, D., Caulk, A. W., Tellides, G., & Humphrey, J. D. (2018). Compromised mechanical homeostasis in arterial aging and associated cardiovascular consequences. Biomechanics and modeling in mechanobiology, 17(5), 1281–1295. https://doi.org/10.1007/s10237-018-1026-7

Fleenor, B. S., Marshall, K. D., Durrant, J. R., Lesniewski, L. A., & Seals, D. R. (2010). Arterial stiffening with ageing is associated with transforming growth factor-β1-related changes in adventitial collagen: reversal by aerobic exercise. The Journal of physiology, 588(20), 3971–3982. https://doi.org/10.1113/jphysiol.2010.194753

Gerhard-Herman, M., Smoot, L. B., Wake, N., Kieran, M. W., Kleinman, M. E., Miller, D. T., … & Gordon, L. B. (2012). Mechanisms of premature vascular aging in children with Hutchinson-Gilford progeria syndrome. Hypertension, 59(1), 92–97. https://doi.org/10.1161/HYPERTENSIONAHA.111.180919

Gordon, L. B., Kleinman, M. E., Miller, D. T., Neuberg, D. S., Giobbie-Hurder, A., Gerhard-Herman, M., … & Fligor, B. (2012). Clinical trial of a farnesyltransferase inhibitor in children with Hutchinson–Gilford progeria syndrome. Proceedings of the National Academy of Sciences, 109(41), 16666–16671. https://doi.org/10.1073/pnas.1202529109

Gordon, L. B., Shappell, H., Massaro, J., D’Agostino, R. B., Brazier, J., Campbell, S. E., … & Kieran, M. W. (2018). Association of lonafarnib treatment vs no treatment with mortality rate in patients with Hutchinson-Gilford progeria syndrome. Jama, 319(16), 1687–1695. doi:10.1001/jama.2018.3264

Hale, C. M., Shrestha, A. L., Khatau, S. B., Stewart-Hutchinson, P. J., Hernandez, L., Stewart, C. L., … & Wirtz, D. (2008). Dysfunctional connections between the nucleus and the actin and microtubule networks in laminopathic models. Biophysical journal, 95(11), 5462–5475. https://doi.org/10.1529/biophysj.108.139428

Humphrey, J. D., Harrison, D. G., Figueroa, C. A., Lacolley, P., & Laurent, S. (2016). Central artery stiffness in hypertension and aging: a problem with cause and consequence. Circulation research, 118(3), 379–381. https://doi.org/10.1161/CIRCRESAHA.115.307722

Humphrey, J. D., Schwartz, M. A., Tellides, G., & Milewicz, D. M. (2015). Role of mechanotransduction in vascular biology: focus on thoracic aortic aneurysms and dissections. Circulation research, 116(8), 1448–1461. https://doi.org/10.1161/CIRCRESAHA.114.304936

Kim, P. H., Luu, J., Heizer, P., Tu, Y., Weston, T. A., Chen, N., … & Hodzic, D. (2018). Disrupting the LINC complex in smooth muscle cells reduces aortic disease in a mouse model of Hutchinson-Gilford progeria syndrome. Science translational medicine, 10(460), eaat7163. DOI: 10.1126/scitranslmed.aat7163

Korneva, A., & Humphrey, J. D. (2018). Maladaptive aortic remodeling in hypertension associates with dysfunctional smooth muscle contractility. American Journal of Physiology-Heart and Circulatory Physiology, 316(2), H265–H278. https://doi.org/10.1152/ajpheart.00503.2017

Korneva, A., Zilberberg, L., Rifkin, D. B., Humphrey, J. D., & Bellini, C. (2019). Absence of LTBP-3 attenuates the aneurysmal phenotype but not spinal effects on the aorta in Marfan syndrome. Biomechanics and modeling in mechanobiology, 18(1), 261–273. https://doi.org/10.1007/s10237-018-1080-1

Kreienkamp, R., Billon, C., Bedia-Diaz, G., Albert, C. J., Toth, Z., Butler, A. A., … & Gonzalo, S. (2019). Doubled lifespan and patient-like pathologies in progeria mice fed high-fat diet. Aging Cell, 18(1), e12852. https://doi.org/10.1111/acel.12852

Laurent, S., & Boutouyrie, P. (2015). The structural factor of hypertension: large and small artery alterations. Circulation research, 116(6), 1007–1021. https://doi.org/10.1161/CIRCRESAHA.116.303596

Lemire, J. M., Patis, C., Gordon, L. B., Sandy, J. D., Toole, B. P., & Weiss, A. S. (2006). Aggrecan expression is substantially and abnormally upregulated in Hutchinson–Gilford progeria syndrome dermal fibroblasts. Mechanisms of ageing and development, 127(8), 660–669. https://doi.org/10.1016/j.mad.2006.03.004

Meredith Jr, J. E., Fazeli, B., & Schwartz, M. A. (1993). The extracellular matrix as a cell survival factor. Molecular biology of the cell, 4(9), 953–961. https://doi.org/10.1091/mbc.4.9.953

Mitchell, G. F., Hwang, S. J., Vasan, R. S., Larson, M. G., Pencina, M. J., Hamburg, N. M., … & Benjamin, E. J. (2010). Arterial stiffness and cardiovascular events: the Framingham Heart Study. Circulation, 121(4), 505. doi: 10.1161/CIRCULATIONAHA.109.886655

Murtada, S. I., Ferruzzi, J., Yanagisawa, H., & Humphrey, J. D. (2016). Reduced biaxial contractility in the descending thoracic aorta of fibulin-5 deficient mice. Journal of biomechanical engineering, 138(5), 051008. https://doi.org/10.1115/1.4032938

Nilsson, P. M., Boutouyrie, P., Cunha, P., Kotsis, V., Narkiewicz, K., Parati, G., … & Laurent, S. (2013). Early vascular ageing in translation: from laboratory investigations to clinical applications in cardiovascular prevention. Journal of hypertension, 31(8), 1517–1526. doi: 10.1097/HJH.0b013e328361e4bd

Olive, M., Harten, I., Mitchell, R., Beers, J. K., Djabali, K., Cao, K., … & Gerhard-Herman, M. (2010). Cardiovascular pathology in Hutchinson-Gilford progeria: correlation with the vascular pathology of aging. Arteriosclerosis, thrombosis, and vascular biology, 30(11), 2301–2309. https://doi.org/10.1161/ATVBAHA.110.209460

Osorio, F. G., Navarro, C. L., Cadiñanos, J., López-Mejía, I. C., Quirós, P. M., Bartoli, C., … & Depetris, D. (2011). Splicing-directed therapy in a new mouse model of human accelerated aging. Science translational medicine, 3(106), 106ra107–106ra107. DOI: 10.1126/scitranslmed.3002847

Prakash, A., Gordon, L. B., Kleinman, M. E., Gurary, E. B., Massaro, J., D’Agostino, R., … & Smoot, L. (2018). Cardiac abnormalities in patients with Hutchinson-Gilford progeria syndrome. JAMA cardiology, 3(4), 326–334. doi:10.1001/jamacardio.2017.5235

Rammos, C., Hendgen-Cotta, U. B., Deenen, R., Pohl, J., Stock, P., Hinzmann, C., … & Rassaf, T. (2014). Age-related vascular gene expression profiling in mice. Mechanisms of ageing and development, 135, 15–23. https://doi.org/10.1016/j.mad.2014.01.001

Roccabianca, S., Bellini, C., & Humphrey, J. D. (2014). Computational modelling suggests good, bad and ugly roles of glycosaminoglycans in arterial wall mechanics and mechanobiology. Journal of The Royal Society Interface, 11(97), 20140397. https://doi.org/10.1098/rsif.2014.0397

Rogers, W. J., Hu, Y. L., Coast, D., Vido, D. A., Kramer, C. M., Pyeritz, R. E., & Reichek, N. (2001). Age-associated changes in regional aortic pulse wave velocity. Journal of the American College of Cardiology, 38(4), 1123–1129. DOI: 10.1016/S0735-1097(01)01504-2

Swift, J., Ivanovska, I. L., Buxboim, A., Harada, T., Dingal, P. D. P., Pinter, J., … & Rehfeldt, F. (2013). Nuclear lamin-A scales with tissue stiffness and enhances matrix-directed differentiation. Science, 341(6149), 1240104. DOI: 10.1126/science.1240104

Valentin, A., Cardamone, L., Baek, S., & Humphrey, J. D. (2008). Complementary vasoactivity and matrix remodelling in arterial adaptations to altered flow and pressure. Journal of The Royal Society Interface, 6(32), 293–306. https://doi.org/10.1098/rsif.2008.0254

Varga, R., Eriksson, M., Erdos, M. R., Olive, M., Harten, I., Kolodgie, F., … & Avallone, H. (2006). Progressive vascular smooth muscle cell defects in a mouse model of Hutchinson–Gilford progeria syndrome. Proceedings of the National Academy of Sciences, 103(9), 3250–3255. https://doi.org/10.1073/pnas.0600012103

Verstraeten, V. L., Ji, J. Y., Cummings, K. S., Lee, R. T., & Lammerding, J. (2008). Increased mechanosensitivity and nuclear stiffness in Hutchinson–Gilford progeria cells: effects of farnesyltransferase inhibitors. Aging cell, 7(3), 383–393. https://doi.org/10.1111/j.1474-9726.2008.00382.x

Villa-Bellosta, R., Rivera-Torres, J., Osorio, F. G., Acín-Pérez, R., Enriquez, J. A., López-Otín, C., & Andrés, V. (2013). Defective extracellular pyrophosphate metabolism promotes vascular calcification in a mouse model of Hutchinson-Gilford progeria syndrome that is ameliorated on pyrophosphate treatment. Circulation, 127(24), 2442–2451. https://doi.org/10.1161/CIRCULATIONAHA.112.000571

Vlachopoulos, C., Aznaouridis, K., & Stefanadis, C. (2010). Prediction of cardiovascular events and all-cause mortality with arterial stiffness: a systematic review and meta-analysis. Journal of the American College of Cardiology, 55(13), 1318–1327. DOI: 10.1016/j.jacc.2009.10.061

Wagenseil, J. E., & Mecham, R. P. (2009). Vascular extracellular matrix and arterial mechanics. Physiological reviews, 89(3), 957–989. https://doi.org/10.1152/physrev.00041.2008

Wheeler, J. B., Mukherjee, R., Stroud, R. E., Jones, J. A., & Ikonomidis, J. S. (2015). Relation of murine thoracic aortic structural and cellular changes with aging to passive and active mechanical properties. Journal of the American Heart Association, 4(3), e001744. https://doi.org/10.1161/JAHA.114.001744

White, L., Haines, H., & Adams, T. (1968). Cardiac output related to body weight in small mammals. Comparative Biochemistry and Physiology, 27, 559–565. https://doi.org/10.1016/0010-406X(68)90252-1

[dataset] Murtada, Sae-Il et al. (2019), Paradoxical Aortic Stiffening and Subsequent Cardiac Dysfunction in Hutchinson-Gilford Progeria Syndrome, v2, Yale, Dataset, https://doi.org/10.5061/dryad.mcvdncjw9

